# DPI-score: A deep learning-based metric for assessing protein-protein interfaces in cryoEM derived assemblies

**DOI:** 10.64898/2026.01.12.699015

**Authors:** Niraj Bhujel, Nicholas Whyatt, Agnel Praveen Joseph, Jyothish Thomas, Luc G. Elliot, Tamilarasan Sabapathy, Jeyan Thiyagalingam, Sony Malhotra

## Abstract

Advances in cryoEM have led to a surge in high-resolution structures; however, model building and refinement at resolutions (>=3 Å) and in regions with variable local resolution remain challenging, making validation essential. CryoEM derived assemblies often contain extensive protein–protein interfaces, however, most existing validation metrics focus on density fit or overall geometry without directly assessing interface quality. To address this, we present DPI-Score, a deep learning-based metric for assessing protein-protein interfaces in cryoEM derived complexes. The method uses only raw structural coordinates of interface atoms, without requiring engineered features, and achieves 87.53% validation accuracy. DPI-Score was applied to 29,120 interfaces from 6,011 fitted entries with resolutions worse than 3 Å in the Electron Microscopy Data Bank. Here, we show that DPI-Score provides complementary information to existing validation metrics and can identify interface errors in modelled assemblies that are not detected by density-based scores alone.

## 1 Introduction

Macromolecular structures can be resolved using various methods such as macromolecular X-ray crystallography, nuclear magnetic resonance, cryogenic-sample electron microscopy (cryoEM[1]). CryoEM has gained popularity due to its ability to image macromolecules in a ‘near-native state’, particularly for large, dynamic macromolecular complexes and those that are difficult to crystallise. Over the years, there have been significant developments in the cryoEM field such as complementary metal-oxide-semiconductor direct imaging detectors for electron microscopy[2], improved sample preparation and image processing methods to decipher high resolution structures[3–5].

Since the ‘resolution revolution’[6], increasingly accessible cryoEM has produced an exponential number of depositions to the Electron Microscopy Data Bank (EMDB)[7] and the Protein Data Bank[8]. Moreover, over the past decade, the number of high resolution entries (*<*=3-4 Å) has increased significantly (105 structures in 2015 vs 7615 structures as of Nov 2025, https://www.ebi.ac.uk/emdb/statistics). However, 23% of the entries in EMDB have resolution worse than 4 Å. Depending on the map resolution, protein(s) can be identified and their model(s) can be built using automated approaches[9]. Additionally, machine learning (ML)-based approaches for generating an initial model, including multi-chain structure prediction through AlphaFold3(AF3)[10] have gained popularity in recent years. Irrespective of the global resolution, this deluge of depositions must be subjected to validation metrics to make sure the errors aren’t propagated downstream for further analysis and interpretation[11, 12].

Resolution in cryoEM is a function of the signal-to-noise ratio[13] that can vary significantly across the map[14]. This variance in local resolution means we cannot rely on one, definitive, global resolution[15] and must assess local model-map fit through comprehensive validation metrics [12]. EMDR (EMDataResource) map and model challenges have compared Fit-to-Map metrics and recommended using multiple scoring methods to obtain full and objective assessment[11]. The commonly used validation approaches using map information include CCC (cross-correlation coefficient[16–18]), EMringer[19], SMOC[20] (segment-based Manders’ overlap coefficient), Q-scores[21]. These metrics assess and guide the fitting of the model to the map at different structural levels and resolution ranges. Additionally, overall model geometry can be assessed using tools like MolProbity[22] and CaBLAM[23], PROCHECK[24], WHATCHECK[25], and Ramachandran Z-scores[26, 27]. Several of these scores can be calculated using the CCP-EM validation task[28] within Doppio(www.ccpem.ac.uk).

However, these validation metrics do not account for potential error at the protein-protein interfaces, which can propagate and may have impact in downstream processes such as function interpretation, drug discovery. When models are built in the cryoEM map, they’re often built sequentially (one chain fitted into the map at a time, independent of the others). Segmentation techniques are not always accurate enough to identify boundaries between subunits. Symmetry options in model building can be ill-applied, or models may have been drawn from maps where the subunit was locally refined. These approaches may lead to suboptimal placement of subunits in the map.

Scores such as the protein-interface quality score (PI-score[29], density independent metric) uses derived interface features such as contact pairs, surface area, shape complementarity, and conserved interface residues to assess the quality of protein-protein interfaces. However, these features are computationally expensive to compute and rely on multiple software to compute these features. There are several published ML methods for predicting protein–protein interactions, which vary in terms of model architecture, training dataset and features used during training. Some recent deep learning-based methods have used interface features to predict the quality of protein-protein interfaces such as DeepRank[30], DeepRank-GNN[31] and DeepRank-GNN-ESM[32]. These methods have been used to score docked poses and to distinguish biological interfaces from crystal contacts, but they have not been specifically applied to the interfaces in cryoEM assemblies. Additionally there have been recent attempts such as AF3Score[33] to provide score-only adaptation of AF3 rather than a structure predictor. It takes an existing biomolecular structure and evaluates its quality using AF3’s internal confidence framework.

In this article, a Deep learning-based Protein Interface score (DPI-score) trained on dynamically sized grid-squares centred on interfaces from high resolution structures, is presented. Considering cryoEM structures are solved at a wide range of resolutions, to simulate interfaces modelled at low resolutions, we added noise to the training dataset by generating structurally close (‘native-like’) docked complexes. It achieves state-of-the art performance on the assessment of protein-protein interfaces, using only the spatial positions of the interface atoms (no map information, complex or derived features needed). The performance of DPI-score to distinguish the ‘near-native’ models was assessed using cryoEM oligomeric targets from CASP15 and CASP16[34] and fitted models obtained from PowerFit[35]. Assessment of the fitted models (resolution worse than 3 Å) associated with the EMDB entries suggests DPI-score can identify the modelling errors at the interface that were undetected by other routinely used density-based metrics. DPI-Score provides a fast and accurate approach for validating protein-protein interfaces and is freely available for academic use on GitLab (https://gitlab.com/ccpem/dpi) or via the CCP-EM software suite, Doppio (https://www.ccpem.ac.uk/software).

## 2 Results

This section presents the DPI-score and its performance across training, bench-marking, and application to independent test datasets. The model is trained on interfaces derived from the Protein Data Bank (PDB)4.1.1) and extensively evaluated on oligomeric cryoEM targets from the CASP15 and CASP16 rounds (https://www.predictioncenter.org/index.cgi) in addition to protein complexes generated by rigid-body fitting using PowerFit[35]. Computational efficiency is assessed by bench-marking training and inference on both CPU and GPU platforms, and the practical utility of DPI-score is demonstrated through representative examples highlighting its ability to identify interface modelling errors in cryoEM derived macromolecular assemblies.

### 2.1 The Dataset

For model training and validation, 6,435 high-resolution complexes (12,825 interfaces; positive dataset 1, PD1) were obtained from the Protein Data Bank[36] using the criteria described in Methods (see section 4.1.1). To enable the CNN to learn features more representative of fitted cryoEM models, PD1 was augmented by introducing small perturbations to the relative positions and orientations of interacting subunits, generating positive dataset 2 (PD2) comprising 4,006 interfaces (see section 4.1.2). A negative dataset (ND) of 9,090 interfaces, structurally distinct from the native interfaces, was also generated (see section 4.1.3). Together, these datasets were partitioned into five stratified folds for cross-validation, with approximately 20,736 interfaces used for training and 5,184 for validation in each fold (see section 4.5 and Table 1).

**Table 1:** Summary of the datasets used for model training, validation, and testing. The dataset derived from the PDB is used for five-fold cross-validation, whereas the PowerFit, CASP15, and CASP16 datasets were used exclusively for independent testing.

| Dataset | Structures | Labelled Interfaces |  |  | Unlabelled Interfaces |
| --- | --- | --- | --- | --- | --- |
|  |  | Positives | Negatives | Total |  |
| PDB | 6,428 | 16,831 | 9,090 | 25,921 | NA |
| PowerFit (Test) | 186 | 1,133 | 74 | 1,207 | 8,811 |
| CASP15 (Test) | 5 | 277 | 41 | 318 | 3,688 |
| CASP16 (Test) | 14 | 3,155 | 116 | 3,271 | 19,451 |

Independent test datasets were created for model evaluation using interfaces derived from oligomeric targets from the two most recent CASP rounds (https://predictioncenter.org) and models obtained using PowerFit[35]. The details on test dataset are described in Methods (see section 4.1.4). The labels for these interfaces were assigned based on their similarity to the target structure. These test interfaces were excluded from model training to enable a stringent evaluation of the model’s generalization performance. Table 1 summarizes all training and test datasets.

### 2.2 Model Training and Evaluation Results

An overview of the 3D CNN model architecture used is shown in Fig.1a. As the input representation includes only the structural coordinates of interface atoms, we used basic 3D CNN architecture in the first version of this method for simplicity, speed, and ease of deployment. This allows us to test the hypothesis if the structural coordinates provide information which can distinguish native/native-like interfaces from a pool of modelled complexes.

The 3D model is trained using the 3D grid representing the (x,y,z) coordinates of interface atoms, without any derived features. To assess the effect of grid representation, we experiment two types of grids : ‘simple’ grid (see Fig. 1b) and ‘dynamic’ grid (see Fig. 1c). A comparison of atomic retention, accuracy and compute time for various grid resolutions (16^3^, 32^3^ and 48^3^) between ‘simple’ and ‘dynamic’ grid representation on our PDB training dataset is shown in Fig. 1d. Dynamic grid representations (see section 4.3 for grid construction details) shows higher atomic retention than ‘simple’ grids, even at smaller grid sizes. Consequently, a compact ‘dynamic’ grid (32^3^) can achieves accuracy comparable to 48^3^ ‘simple’ grid, with substantially lower compute time. Moreover, ‘dynamic grid’ based representation improves training behaviour, achieving faster convergence and lower training loss than ‘simple grid’ representation, as shown in Fig. 1e.

**Fig. 1:**
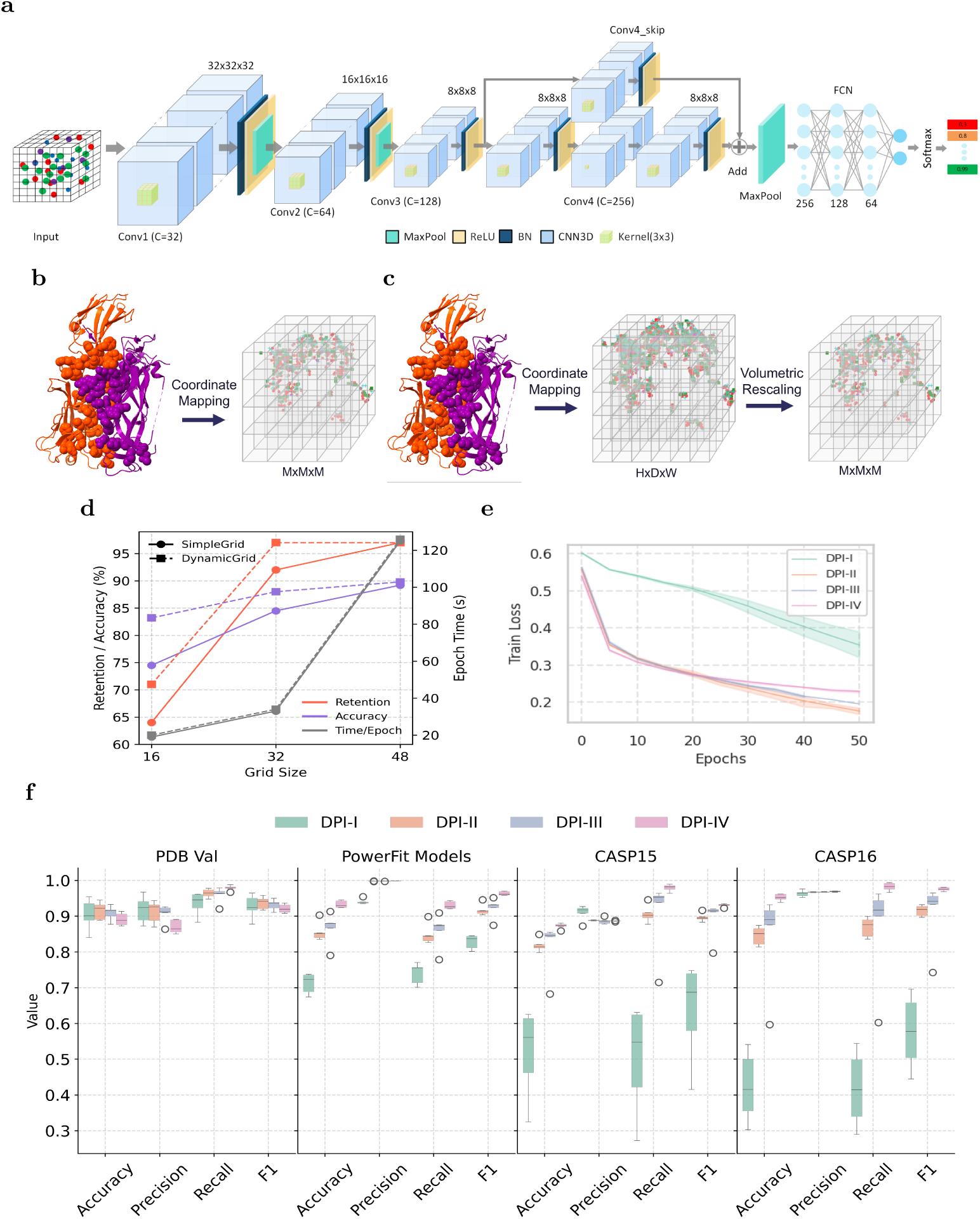
DPI evaluation results on validation and test datasets. **a.** CNN model architecture. **b.** Simple-grid representation with atom coordinates mapped to a pre-defined 3D voxel grid. **c.** Dynamic grid representations, where each atom coordinate is mapped to a voxel of size 1 Å^3^ and then volumetrically rescaled to a target grid shape. **d.** Accuracy, retention ratio, and computation time of simple and dynamic grid representations for different grid sizes. **e.** Training behaviour of ‘simple’ and ‘dynamic’ grid-based models. **f.** Performance metrics of DPI models on validation (first panel) and test datasets (second-fourth panels). Each bar’s centre line shows the median, the box edges the 25th–75th percentiles, and whiskers are set to 1.5 times the interquartile range.

Four variants of inputs representations were experimented to systematically assess the contribution from each component to the prediction performance.

- DPI-I: Simple grid representations
- DPI-II: Dynamic grid representations
- DPI-III: Dynamic grid with additional channel for steric clash
- DPI-IV: Dynamic grid with steric clash channel and data augmentation

The first two variants DPI-I and DPI-II are explained in section 4.3. DPI-III refers to dynamic grid with additional channel for steric clashes that is defined as instances where the Euclidean distance between two atomic centres falls below the sum of their respective van der Waals radii. Additionally, data augmentation methods such as random rotations and scaling (see section 4.6) were applied to DPI-III during training for improving model robustness to unknown orientations in DPI-IV.

All DPI model variants were trained and evaluated using K-fold cross-validation (see section 4.5). For each fold and DPI variant, performance metrics; accuracy, precision, recall and F1 scores (see section 4.9) were calculated on the corresponding validation fold, and averaged across all folds. Fig. 1f (first panel) shows the mean and median of these metrics. The simple grid representation of the input data (DPI-I) provides an strong baseline performance demonstrating the effectiveness of learning using only the structural coordinates. The dynamic grid approach (DPI-II) has a narrower distribution as compared to the simple grid approach (DPI-I) indicating the robustness likely resulting from higher atomic retention of the dynamic grid representations. Model robustness and generalization can be further improved by adding steric clash information to the dynamic grids (DPI-III). Random rotation of the grids during training (DPI-IV) only improves recall at the cost of other metrics however it generalise better on the unseen test datasets (see 2.3 for more details on performance on test dataset).

The confusion matrices for each model variant across all datasets are provided in Supplementary Fig. 1 for a more detailed comparison. An evaluation of four variants using both Receiver Operating Characteristic (ROC) and Precision-Recall (PR) curves across five cross-validation folds is shown in Supplementary Fig. 2. Based on these comparisons, DPI-IV was used as the final model for comparative analysis throughout the paper due to its better performance and stability.

### 2.3 Performance on Test Datasets

The following datasets were used to assess the performance of the DPI-score (see Table 1 for more details):

1. CryoEM oligomeric targets from the two most recent rounds of CASP (https://predictioncenter.org) experiments - CASP15 and CASP16.
2. Models obtained using a rigid body fitting method - PowerFit[35]

Five and fourteen oligomeric cryoEM targets from CASP15 and CASP16, respectively, were assessed using DPI-score (see Supplementary Table 1). The computed DPI-scores for the CASP models are provided as Supplementary Table 2. The confusion matrices for all model variants on all test datasets are provided in Supplementary Fig. 1, the ROC curves and the precision-recall (PR) curves are shown in Fig. 2a and Supplementary Fig. 3, respectively. These reveals a much more challenging regime for CASP15 and CASP16 interfaces with AUC values of 0.61 and 0.58 respectively. In both cases, ROC curve lie above the random baseline, demonstrating non-random ranking capability, however, the proximity of the ROC curve to the diagonal and the modest AUC value indicates substantial overlap between positive and negative score distributions, reflecting the intrinsic difficulty in CASP datasets, which is explained below. Other performance metrics (Accuracy, Precision, Recall and F1) for different model variants on the test datasets (PowerFit, CASP15, and CASP16) are shown in the Fig. 1f. Although simple grid representations demonstrated competitive performance in the validation dataset, it could not generalise well for the test datasets (Fig. 1f). Consistent with the performance on validation dataset and during training, DPI-III (dynamic grid with additional steric clash information) and DPI-IV (with data augmentation) improve most of the metrics during testing, except for the precision on CASP15 models. We believe this may be due to:

1. High false positive rates for some targets such as H1129 (see Supplementary Fig. 4).
2. Significant fractions of interfaces were not assigned labels in all the three datasets (Table 1)-CASP15 (92%), CASP16 (43%) and PowerFit (88%) and the number of negatively labelled interfaces is also very low (CASP15-∼1% and CASP16-∼ 1.7%).

**Fig. 2:**
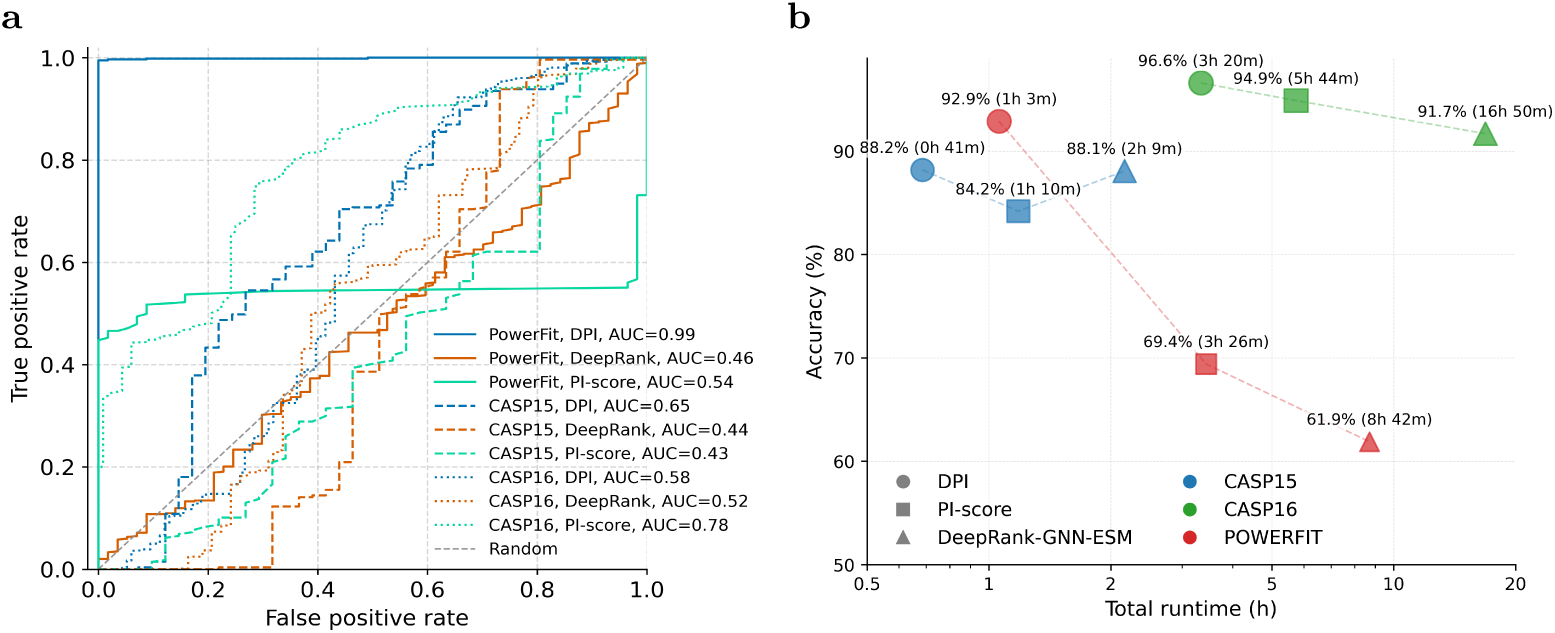
Comparisons with existing methods. **a.** Comparison of ROC-AUC curve on test datasets. **b.** Comparison of accuracy and computational runtime. All models were run on same hardware with Intel XG 5218 CPU@2.30 GHz and NVIDIA Tesla V100 GPU. Runtime denotes the total computational time required for loading the interface, processing, and score prediction. All metrics are computed on a subset of interfaces common to all models.

The IS-score vs iRMSD distribution for the labelled interfaces of CASP15 and CASP16 (Supplementary Figs. 4 and 5 respectively) suggest clusters of true positives (salmon) and true negatives (green) for positives and negative interfaces respectively. Unlabelled interfaces (the majority of the targets) with high iRMSD and low IS-score were predicted to have lower DPI-score (class0, in green), whereas the interfaces closer to the interface in the target structure i.e. low iRMSD and high IS-score were assigned to positive class (class1, in pink) (Supplementary Figs. 6).

There were some targets (`H1236, T1234o, T1235o` and `T1249v2o` from CASP16) with a high number of unlabelled positives (Supplementary Fig. 6a., pink in the region with high iRMSD and low IS-score). It is interesting to note that the three out of these four targets are linked to the same PDB ID: 8QPQ; H1236 is the whole complex (A6B6C3), `T1234o` is the C3 (prokaryotic polysaccharide deacetylase trimer) subcomplex and `T1235o` is the A6 (hexamer of gp30) subcomplex. Although the unlabelled interfaces could not meet the (strict) threshold criteria for both IS-score and iRMSD, the majority of the interfaces have iRMSD *<* 3 Å, implying they may be structurally close to the target interface. IS-score takes conservation of the interface contacts into account, therefore it is possible that interfaces in these ‘false positives’ are slightly shifted relative to the native interface (low iRMSD, yet low IS-score).

The labelled and unlabelled interfaces from the PowerFit generated models follow the expected trend *i.e.* the interfaces with low IS-score and high iRMSD obtain a low DPI-score (green, Supplementary Fig. 7 and high scoring (DPI-score) interfaces have high IS-score and low iRMSD (pink, Supplementary Fig. 7).

### 2.4 Benchmarking DPI-score

Table 2 presents the average processing time per interface across different stages of training and inference. Feature extraction which includes obtaining coordinates of interface atoms, is performed once and remains constant across all input representations, whereas grid generation times vary according to the grid types and features used. The reported measurements exclude data loading times, as these are highly dependent on storage hardware and system I/O capabilities. The feature extraction and grid generation were executed on CPUs, while training and inference were performed on both GPUs and the CPUs. The total training time per model varied due to the use of early stopping, which resulted in different numbers of training epochs (see section 4.4 for more details). All four DPI-score variants showed comparable inference speeds, requiring 1.7-1.8 ms per complex on a V100 GPU and 68.1-89.0 ms on CPU. DPI-II training the fastest (19.4 min) and DPI-IV the slowest (84.5 min).

**Table 2:** Average speed per interface measured on Intel Xeon Gold 5218 CPU at 2.30 GHz(28 cores, 28 threads) and NVIDIA Tesla V100 GPU. Lower values indicate higher throughput.

| Model | Dataset | Feature extraction (ms) | Grid generation (ms) | Training time (epochs) | Inference (ms) |  |
| --- | --- | --- | --- | --- | --- | --- |
|  |  |  |  | V100 | V100 | CPU |
| DPI-I | PDB | 29.6 | 65.8 | 25.6min (69 epochs) | 1.7 | 73.3 |
| DPI-II |  |  | 106.1 | 19.4mins (54 epochs) | 1.8 | 68.1 |
| DPI-III |  |  | 109.7 | 40.6mins (54 epochs) | 1.8 | 77.6 |
| DPI-IV |  |  | 109.7 | 84.5mins (69 epochs) | 1.8 | 89.0 |

The performance of DPI-score (DPI-IV) was compared to two other machine learning-based methods, namely PI-score[29] and DeepRank-GNN-ESM[32] using the test datasets (Table 1). PI-score uses conservation as one of interface features, but this is a very time-consuming step as it requires identification of homologs and computing their alignment which is used as an input to an algorithm (rate4site) to compute conservation scores. Therefore, the conservation free version of PI-score (https://gitlab.com/topf-lab/pi_score) was used. Similarly, DeepRank-GNN requires position specific scoring matrices (PSSM) as a required input, hence the most recent version which is PSSM-free, DeepRank-GNN-ESM, was used to score all the interfaces in the validation set. It is to be noted that DeepRank-GNN-ESM is a method for scoring protein-protein models however is not specifically trained to assess interfaces in cryoEM derived assemblies.

These comparisons are presented as Fig. 2 for a set of common labelled interfaces from the test datasets which were successfully processed and scored by all three methods. DPI-score achieved the highest accuracy across all three test datasets, with the largest performance gain observed on the models fitted using PowerFit(Fig. 2b).

It’s worth highlighting that these other two methods (PI-score and DeepRank-GNN-ESM) rely on a number of interface parameters and take significantly longer than DPI-score (approximately 26 hours and 9 hours in total for DeepRank-GNN-ESM and PI-score, respectively, for the entire dataset). On the other hand, DPI-score took only about 4 hours. The mean average time (in sec) to score per interface was 3.8, 20.5 and 7.6 for DPI-score, DeepRank-GNN-ESM and PI-score, respectively.

Fig. 2a shows the ROC curves of all three models on test datasets, where ROC curve of DPI was obtained by averaging scores across all five-folds. DPI-score achieves very strong performance on PowerFit dataset (AUC=0.99), substantially outperforming DeepRank-GNN-ESM (AUC=0.46) and PI-score (AUC=0.54). On the CASP datasets, performance differences are less clear. On the CASP16 dataset, PI-score achieved a higher ROC-AUC (0.78) than DPI-score (0.58). One possible explanation is that PI-score benefits from explicitly engineered interface descriptors, whereas DPI-score relies solely on learning discriminative representations from atomic coordinates. The challenging nature of the CASP16 targets may favour the use of these pre-computed structural features. Future work will investigate whether more sophisticated model architectures further improve performance on such difficult targets. For the CASP15 dataset, DPI-score achieved AUC of 0.65 while PI-score attains AUC of 0.43.

It should be noted that the methods tested here differ in terms of their model architectures, input representations and hence are best interpreted as operational benchmarking comparisons rather than direct performance comparisons between different model architectures.

### 2.5 Examples from Test Datasets

Below are few examples from the test dataset that demonstrate the performance and application of DPI-score.

#### 2.5.1 CASP15 target: T1121o

For the target `T1121o`, 3.7 Å resolution homodimer from *Pseudomonas Aeruginosa*, 79 interfaces were assessed. The interface in the target structure obtains a high DPI-score (Fig. 3a). Four (`TS098_4o, TS098_5o, TS165_2o` and `TS234_1o`) of these interfaces obtained low DPI-score (*<*0.5) which are shown in Fig. 3b-e. For these models the iRMSD and IS-scores imply that they are far from the target structure (Supplementary Table 2).

**Fig. 3:**
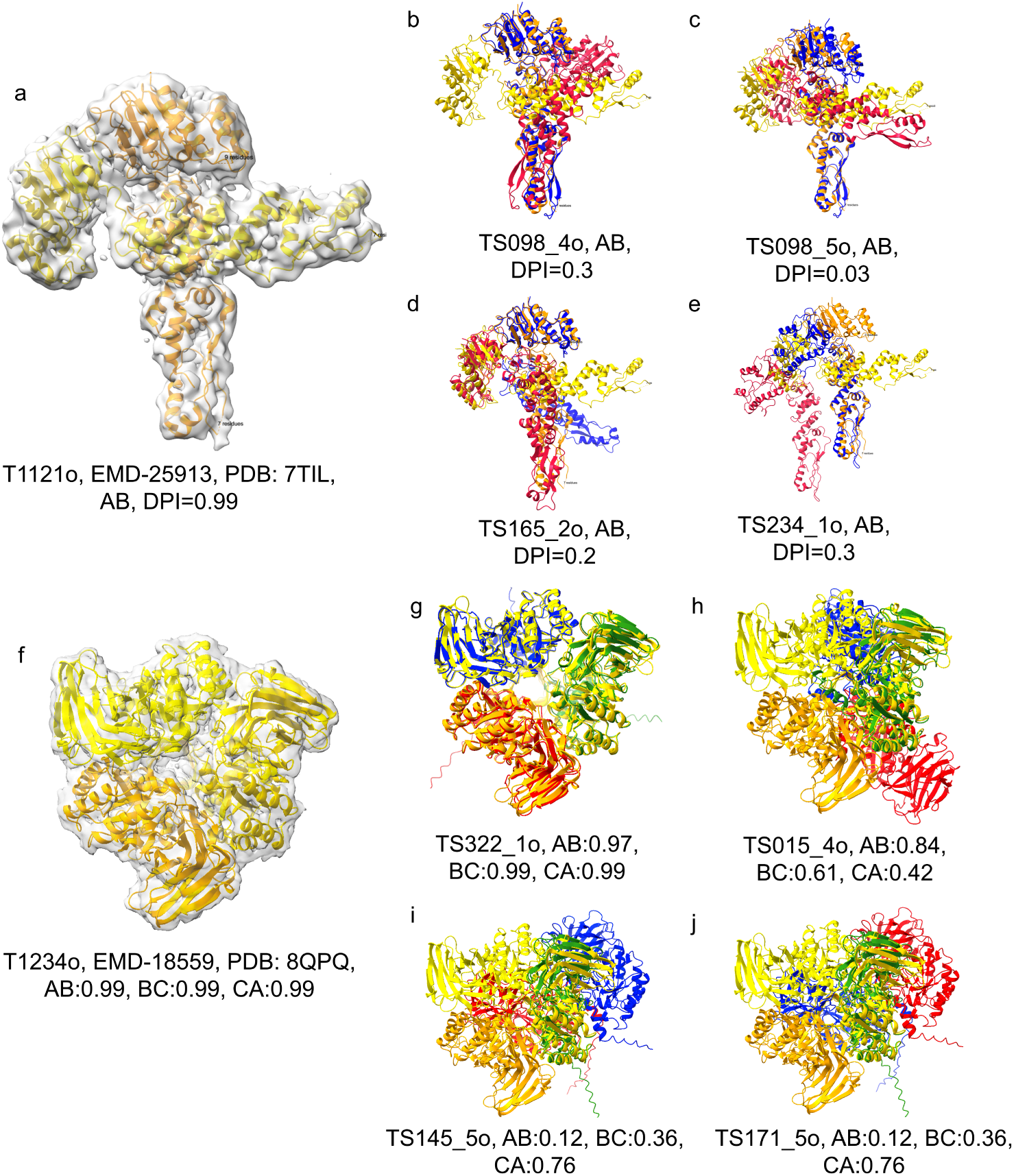
Scoring the interfaces in the oligomeric CASP15 target `T1121o` and CASP16 target `T1234o`. In all the panels, chains from the target structure are shown in orange and yellow and the chains from assessed model are shown in red, blue and green. a. Target structure (PDB: 7TIL) for `T1121o` within its map (EMDB: 25913). b-e. Interfaces in the models `TS098_4o, TS098_5o, TS165_2o` and `TS234_1o`, respectively, for the target `T1121o` with their DPI-scores. f. Target structure (PDB: 8QPQ) for `T1234o` within its map (EMDB: 18559). g. Interfaces from the model `TS322` 1o for target `T1234o`, were scored high, and h-j. Interfaces in the models `TS015_4o, TS145_5o` and `TS171_5o` respectively, were scored low using DPI-score.

To demonstrate the ability of DPI-score to distinguish ‘target-like’ interfaces, for all these four models (interfaces), we used RELION[37] (image-handler, low pass utility) to generate lower resolution density maps at 5, 8, 10 and 12 Å. As the resolution decreases, it becomes more difficult to distinguish poorly modelled interfaces using density-based metrics such as the CCC(see Supplementary Table 3). This is particularly evident in the case of model `TS098_5o`, with high CCC score (0.82) at 12 Å despite having a low DPI-score (0.03).

#### 2.5.2 CASP16 target: T1234o

991 interfaces (from 329 models) were assessed for the target `T1234o`, which is a prokaryotic polysaccharide deacetylase homotrimer solved at 2.7 Å global resolution within the complex structure of full Haloferax tailed virus 1 (Fig. 3f). All the interfaces (between chains AB, BC and CA) in the target structure obtain high DPI-scores (Fig. 3f). The interfaces from selected models with high and low scoring interfaces are shown in Fig. 3 (g-j). All the three interfaces in model `TS322_1o` (see Fig. 3g) are very close to the interfaces in target structure and rightly obtain high DPI-scores. For the three other models `TS015_4o, TS145_5o` and `TS171_5o`, all or some of the interfaces scored low on DPI-scores and it can be clearly seen in Fig. 3h-j that they are far from the interfaces observed in the target structure.

#### 2.5.3 Fitted models obtained using PowerFit

Rigid fitting of subunit or domain models is a common way to build atomic model of an assembly in a cryoEM map, especially since the development of AlphaFold. Methods such as PowerFit[35] propose a set of solutions which are usually ranked using density dependent metrics such as CCC, which evaluate the quality of fit of the model in a given cryoEM density map. However, these density-based metrics do not reflect the quality of the model and certainly do not assess the quality of the protein-protein interfaces in the modelled complex.

At low resolutions multiple solutions might fit the map with the similar CCC which makes it difficult to identify the true solution from the set[38]. Fig. 4 demonstrates examples where interface focused scores such as DPI-score can help identify errors at the protein-protein interface, which could not be identified using density-based scores such as CCC. We used PowerFit to generate fitted poses for selected chains in maps of resolutions worse than 3 Å and assessed the interfaces of best scoring solutions. To simulate low-resolution fitting, we blurred the map further using Gaussian convolution (see section 4.1.4). We used the deposited model as reference to compare the fitted solutions.

**Fig. 4:**
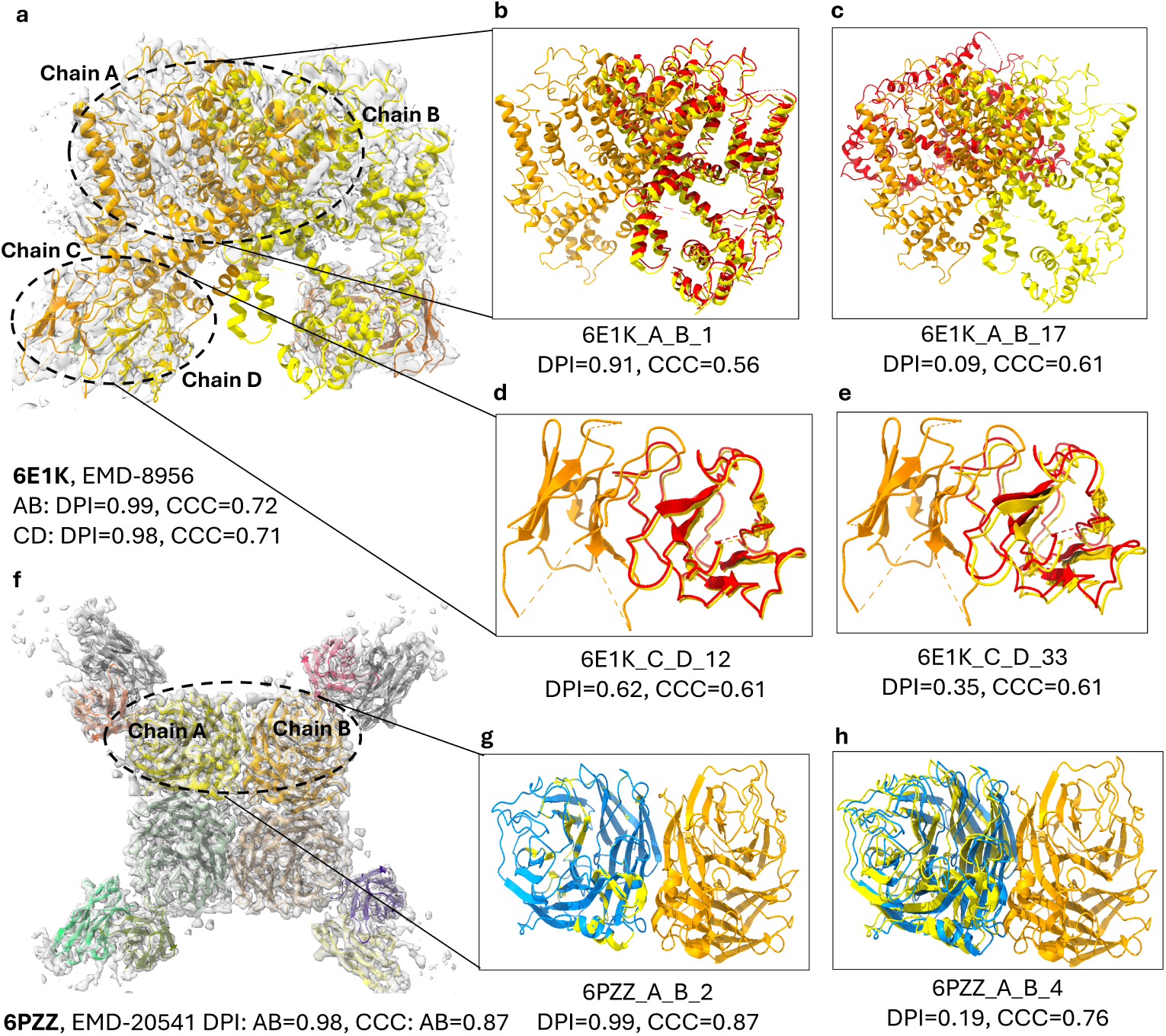
Scoring the interfaces in the fitted models obtained using PowerFit. In all the panels, chains from the target structure are shown in orange and yellow. a. Target structure (PDB ID: 6E1K) within its map (EMDB: 8956). b and c. High and low DPI-scoring models `6E1K_A_B1` and `6E1K_A_B_17`, for interface between chains A and B, shown in red. d and e. High and low DPI-scoring models `6E1K_C_D_12` and `6E1K_C_D_33`, for interface between chains C and D, f. Target structure (PDB ID: 6PZZ) within its map (EMDB: 20541). g and h. High and low DPI-scoring models `6PZZ_A_B_2` and `6E1K_A_B_4`, for the interface between chains A and B.

Calcium channel protein: The interfaces between the channel protein subunits (chains A and B) and between the light and heavy chain of the antibody (chains C and D) in the 3.3 Å structure of calcium channel protein bound to antibodies (EMD-8956, PDBID: 6E1K) obtain high DPI-scores (see Fig. 4**a**), and the interface chains AB and AC have a CCC of 0.72 and 0.71, respectively. We compared the fitted solutions obtained using PowerFit against the deposited model and scored them using DPI-score. Fig. 4**b** shows one of the fitted solutions (`6E1K_A_B_1`) for the calcium channel protein (chains A and B) which is close to the target structure and obtains a high DPIscore of 0.91 and CCC score of 0.56. Another fitted solution (`6E1K_A_B_17`, Fig. 4**c**) for chains A and B clearly has a wrong orientation of the interface (DPI score=0.09), however obtains a CCC score of 0.61 (better than the model with interface structurally close to target structure (`6E1K_A_B_1`)). This highlights the importance of using interface-based scores along with density-based scores such as CCC in order to filter out the correct fitting solutions. For the interfaces between the light and heavy chains of antibody (chains C and D) in the same structure, the CCC score is the same between the fitted models (`6E1K_C_D_12` and `6E1K_C_D_33`) however they can be distinguished using the DPI-score (Fig. 4**d** and Fig. 4**e**).

Neuraminidase: Another case highlighted is the structure of neuraminidase (3.6 Å) which forms a homotetramer (EMDB-20541 and PDB ID: 6PZZ, Fig. 4**f**). The interface from the model deposited in the PDB (‘target’) obtains a high DPI-score (0.98) and the CCC for the two chains (A and B) forming the interface is 0.87. One of the fitting solutions which is structurally very close to the interface in the target obtains a high DPI-score(0.99) and CCC score of 0.87 (same as the chains in target structure, Fig. 4**g**). Another fitting solution which has an offset at the interface (chains A and B) and is recognised by DPI-score(0.19) obtains the CCC score of 0.76 (Fig. 4**h**). This shows that DPI-score is sensitive to discriminate interface quality which is less straightforward with CCC especially at such resolutions.

These cases highlight the importance of using an additional interface centric metric while assessing the quality of fitted models in the cryoEM map when they can’t be distinguished easily using the density-based scores such as CCC.

### 2.6 Application to fitted entries in EMDB

EMDB entries with fitted models with a resolution worse than 3 Å(6010, as of October 2024) were scored using the CCP-EM validation suite[39] to calculate a number of validation metrics, namely Clashscore, MolProbity score, SMOC Z-score *<* −2, average SMOC, average model-map FSC (0.5 cutoff), CCC, CCC mask and overlap score. These scores assess model stereochemistry and also fit-to-map. The valid interfaces (29k) from these entries were scored using DPI-score (see Supplementary Table 4).

DPI-score was observed to have very low correlation to most of the density-based scores Supplementary Fig. 8, which suggests that the subunit interface quality is not directly related to the goodness of model fit. The following examples highlight the specific applications of DPI-score, where the density-based scores alone are not reflective of the quality of the modelled interfaces.

#### 2.6.1 Actomyosin complex in skeletal sarcomere

This is a 6.6 Å resolution structure with a homo pentamer of Actin (EMDB: 13991, PDB ID: 7QIN, Fig. 5**a**). Interfaces between the actin subunits-chains C and E and chains A and C obtain a low DPI-score of 0.19 and 0.43 whereas the density-based scores ccc-mask and average model-map FSC0.5 for the deposited model (PDB ID: 7QIN) are 0.54 (percentile: 49) and 0.76 (percentile: 65) respectively. The averaged iSMOC score for interface residues is 0.92 for both the AC and CE interfaces. The map fit scores at or above 50th percentile suggests that the model compares well (better than 50%) against other models built in maps at similar resolutions. The low DPI scores however suggest that these interfaces may not be optimally modelled in the deposited structure. To check whether the interface could be improved by further automated refinement, CCP-EM software suite v2 (Doppio) was used to perform flexible fitting using TEMPy-REFF[40] job, which was followed by Servalcat refinement[41]. Upon re-refinement, the clashes at the interfaces were resolved which resulted in the overall improvement as indicated by the better density-based scores (CCC-mask=0.6; percentile 57.9, average model-mask FSC0.5=0.89; percentile 98.5), local residue quality scores - SMOC, and the interface scores, DPI-score (for chains C and E= 0.98 and for chains A and C = 0.97). For both the deposited and the refined models, the plots for the SMOC score (per residue) are shown in Fig. 5**a**, for the three interface forming chains. The SMOC scores are overall better for the re-refined structures and interface residues (red circles) are observed to score better, especially for chain E. iSMOC scores upon refinement improve to 0.94 for AC interface and remains unchanged (0.92) for the CE interface Fig. 5**a**.

**Fig. 5:**
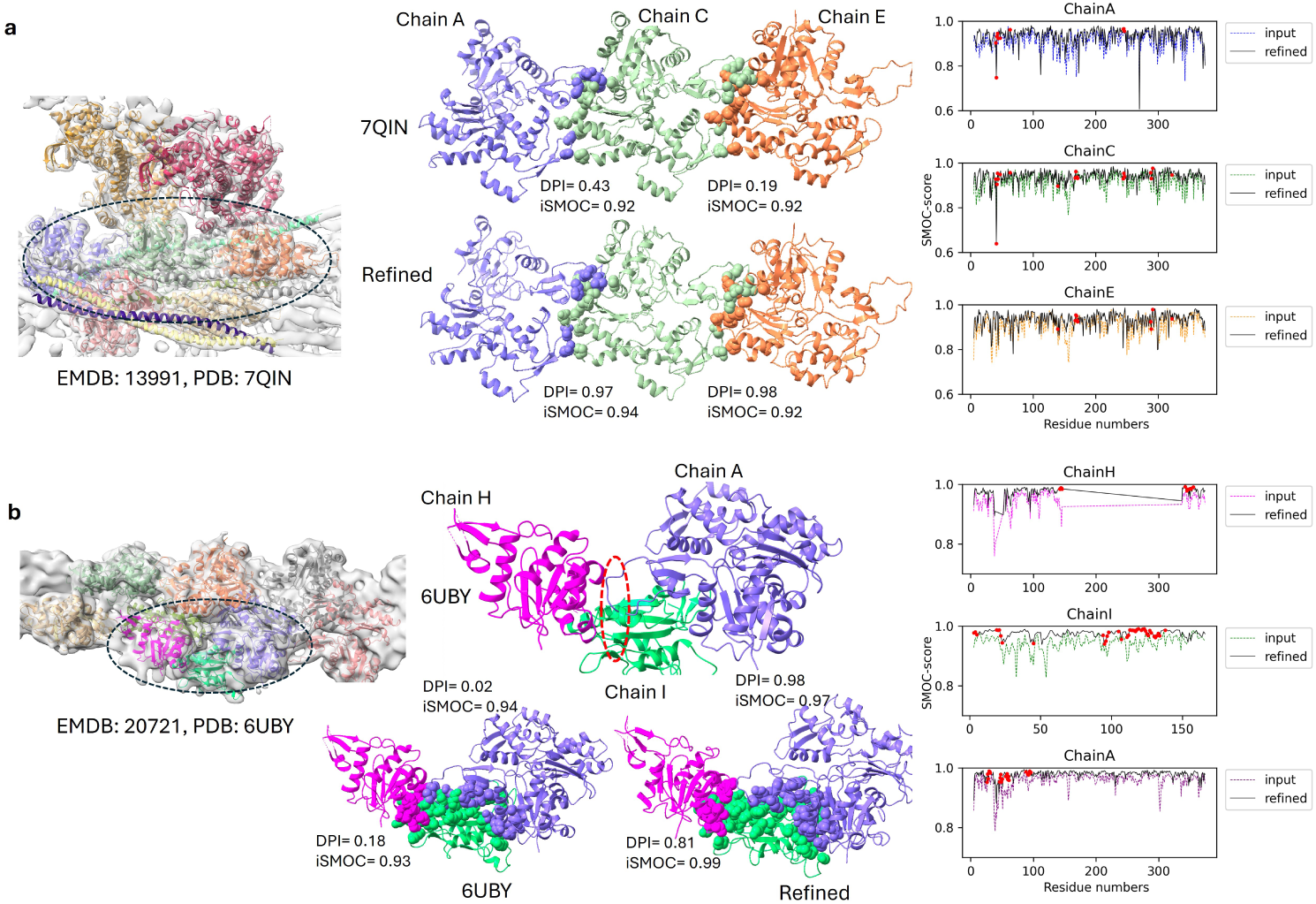
Application to the fitted models in EMDB. The atoms of the interface residues from the chains are shown as spheres for both the fitted model deposited in EMDB and the re-refined model. The plot of the local density-based score (SMOC) is shown for the chains forming an interface in the model and re-refined structures. The interface residues are shown as red spheres on SMOC score plots. a. 6.6 Å resolution structure of actomyosin complex in the skeletal sarcomere (EMD-13991; fitted PDB: 7QIN). The black dashed oval represents the three chains (A, C and E) forming the subcomplex that was investigated for the quality of interfaces. b. 7.5 Å resolution structure of the cofilin-actin complex (EMD-20721, fitted PDB: 6UBY). The black dashed oval represents the three chains (H, I and A) forming the subcomplex that was investigated for the quality of interfaces. The red oval highlights the knotted loop from chain A in chain I.

#### 2.6.2 Cofilin-Actin complex

It is a 7.5 Å structure of the cofilin-actin complex (EMDB: 20721, PDB ID: 6UBY, Fig. 5**b**). Cofilin (chain I) interacts with two actin chains (chains A and H). The interfaces between chains A and I and H and I have low DPI-scores of 0.02 and 0.18, respectively, implying that the interfaces may be sub-optimal. However, the density-based scores for the deposited model are CCC-mask=0.68 (percentile= 75.4) and average model-mask FSC0.5=0.72 (percentile= 44.9). The iSMOC score over the HI and AI interfaces is 0.93 and 0.94, respectively. There is a severe steric clash where one of the loops from the actin chain (H) is knotted (Fig. 6b, shown in red dashed oval) within one of the *α*-helices of the cofilin chain (I). To resolve this severe clash, the knotted region (5 residues, residues 45-49) in the loop was manually translated (unknotted) in UCSF Chimera[42] and further re-refined using TEMPy-REFF followed by Servalcat (Fig. 5b). Upon re-refinement, DPI-scores improve to 0.98 and 0.81 for interfaces between chains A and I and chains H and I, respectively. There is also an improvement in the global density-based scores (ccc-mask=0.81; percentile= 97.2 and average model-mask FSC0.5=0.87; percentile= 96.8) and iSMOC improves to 0.99(HI) and 0.97(AI). The local density quality score-SMOC (per-residue) was also observed to be better for all three chains upon re-refinement Fig. 5**b**, interface residues shown as red circles). This demonstrates the application of interface-based scores such as DPI-score which can help identify sub-optimal protein-protein interfaces in modelled complexes especially in cases where density-based scores alone do not clearly highlight such specific local issues.

## 3 Discussion

### 3.1 Protein Interface Quality Assessment Metric: DPI-score

There are several machine learning methods that assess the quality of protein-protein interfaces using protein sequence and structure features[43], and are routinely used for predicting protein binding sites. Recent methods such as DeepRank[44], DeepRank-GNN[31] and DeepRank-GNN-ESM[32] have applied feature-based classifiers to score docked poses and to distinguish biological contacts from crystal contacts. In 2021, we proposed an interface focused (density-independent) metric, PI-score, to detect errors at the protein-protein interfaces, and demonstrated its application to the models derived using cryoEM[29]. In this work, a deep learning-based metric (DPI-score) to assess the quality of protein-protein interfaces using only structural coordinates of the interface atoms is presented. The training dataset comprises of protein-protein interfaces derived from the high resolution structures determined using either X-ray crystallography or cryoEM (PD1). This positive dataset was further supplemented to add structurally close interfaces obtained by slightly perturbing the relative positions and orientations of the interacting subunits using protein-protein docking (PD2). A negative dataset was obtained from interfaces which were structurally far from the target interface (generated using protein-protein docking) in PD1. DPI-score calculation is computationally efficient, with an average inference time of approximately 1–2 ms per interface (Table 2) with no derived features required as an input into the deep learning model.

In order to effectively encode the atomic coordinates for wider range of interfaces sizes, dynamic volumetric grid representations were used, which outperformed the approach of using a fixed grid size (simple grid). The CNN model inherently learns the relative positions of interface atoms to discriminate between the positive and negative class labels. The presence of steric clashes was penalised during the training. Based on the performance of DPI-score on multiple test datasets, dynamic grid approach was observed to eliminate the trade-off between information preservation and computational efficiency by guaranteeing that most atomic coordinates are captured in the initial representation before standardisation, thereby preventing the truncation artifacts inherent in simple grids.

DPI-score is both accurate (87.54% validation accuracy) and computationally efficient. It is 7.5 times faster than PI-score[29] and 3.4 fold faster than DeepRank-GNN-ESM[32], which achieved validation accuracies of 84% and 60%, respectively.

### 3.2 Applications of DPI-score

The specific applications of DPI-score are discussed:

1. To distinguish near-native models from a pool of modelled solutions (oligomeric cryoEM targets from CASP rounds and fitted solutions obtained using rigid body fitting methods).
2. To validate and detect errors at the protein-protein interfaces in the fitted models associated with EMDB entries.

Approximately 95% of cryoEM structures contain two or more protein chains, making protein-protein interfaces highly prevalent. Several factors can contribute to sub-optimal modelling of these interfaces in the complex structures especially at low-to-intermediate resolution. Challenges such as lower resolution (less than 3 Å and variable local resolution can significantly impact the accuracy of the modelled protein-protein interactions. Interface focused scores such as PI-score[29] and DPI-score can be used to validate and to aid the modelling of interfaces in cryoEM derived assemblies.

CASP style experiments provide an ideal test-bed for evaluating and validating scores such as DPI-score. In these experiments, participating teams model protein complexes without access to the corresponding cryoEM maps. Additionally, the experimentally determined target structures are made available, serving as a ‘ground truth’/reference for assessing the accuracy of the predicted models. Oligomeric cryoEM targets from the two most recent CASP rounds-CASP15 and CASP16 were used to assess the quality of DPI-score and modelled interfaces. For the majority of the targets, DPI-score was effective in distinguishing near-native interfaces from the pool of modelled interfaces. However, some targets - `H1137, T1234o` and `H1236` - were challenging cases from CASP15 and CASP16 where interfaces with low IS-score obtained high DPI-score, however they do have low iRMSD(*<*= 3 Å). IS-score explicitly takes conservation of interface contacts into account, whereas iRMSD is a purely geometric measure of how closely interface atoms align after superposition. Therefore, it is possible that interfaces in these targets are slightly shifted relative to the native interface, resulting in low iRMSD, yet the native contacts are not preserved leading to a low IS-score. We also speculate that for these challenging targets, the spatial positioning of atoms alone does not provide sufficient information to reliably distinguish near-native models and a graph-based approach (interface contact information), such as using Graph Neural Networks (GNNs), may be more suitable than traditional CNNs.

Another application demonstrated is DPI-score’s ability to evaluate fitted solutions obtained through rigid-body fitting method (e.g. PowerFit). In this scenario, the cryoEM map is available during modelling, allowing direct fitting of subunits into the density. However, the assessment is often heavily biased toward density-based metrics, such as CCC, which are not specifically designed to detect errors at the interfaces. By combining density-based scores with interface-focused scores (such as DPI-score) and other standard model validation metrics, a more comprehensive and reliable assessment of the fitted structures can be achieved.

Additional applications of DPI-score include application to validate the modelled complexes obtained using AF3[45]. The observed positive correlation between the ipTM score from AF3 and DPI-score[45] suggests that DPI-score can serve as a reliable orthogonal metric for model validation. It would therefore be of interest to systematically quantify the correlation between these two scores across a broad set of AF3 modelled complexes.

### 3.3 Comparison with Density-based Scores: Application to Fitted Models in EMDB

With the advancement in the field of cryoEM, reconstructions obtained using single-particle analysis are getting close to near-atomic resolution, however only 30% of the entries in EMDB have resolution less than 3 Å (https://www.ebi.ac.uk/emdb/statistics/emdb_resolution_year). Additionally, local resolution within a map can vary significantly.

For cryoEM map interpretation, subunit or domain models are often fitted sequentially followed by refinement of the fitted models. The inter-atom contacts are not usually optimised explicitly in a standard refinement process. If the information in the map is sufficient to position the atoms reliably and the model stereochemistry is optimal, the final model is expected to be of good quality. However, lack of clear validation references for a well-fitted model given the map, often results in many deposited models being under-refined[46]. Also, it is much harder to build models in low-resolution areas of the map without the availability of sufficient additional external restraints or priors.

Various metrics are routinely used to assess the quality of modelled assemblies in cryoEM maps. The global density-based scores highlight the overall quality of fit-to-data of the model and local scores are focused on specific parts of the structure such as per-atom, per residue scores etc. The correlation of DPI-score with other model quality scores such as MolProbity[22] and other density-based scores is very low, indicating that it is a complementary assessment metric. The local scores such as SMOC (per-residue scores), when averaged over interface residues were not reflective of interface quality.

Therefore, DPI-score which is a density-independent metric, when used in combination with other validation metrics (as it provides complementary information) can help identify errors at protein-protein interfaces especially at low-to-intermediate resolution. When applied to fitted models associated with EMDB entries (resolution worse than 3 Å), DPI-score highlights cases in which density-based scores alone fail to identify errors at the interface. Re-refinement of these submitted models leads to improvement in both interface score (DPI-score) density-based metrics.

### 3.4 Availability and Future Directions

The Python-based inference codes along with the trained model weights for DPI-score are available on GitLab at https://gitlab.com/ccpem/dpi. We also provide the GUI version in the CCP-EM software suite Doppio at https://www.ccpem.ac.uk/software for the model validation job (see Supplementary Fig. 9). In future, different model architectures (such as E3NN[47]) and other machine learning approaches such as single label semi-supervised classification will be explored. DPI-score will be a crucial addition to the currently used validation scores.

## 4 Methods

### 4.1 Dataset Description

#### 4.1.1 Dataset of high-resolution complexes (Positive Dataset 1-PD1, ‘native’ interfaces)

High resolution structures resolved through X-Ray and cryoEM, were obtained from PDB[8]. The query was performed on the 3rd of April 2024 for non-redundant representatives at 50% sequence identity, under the criteria laid out in Table 3.

**Table 3:**
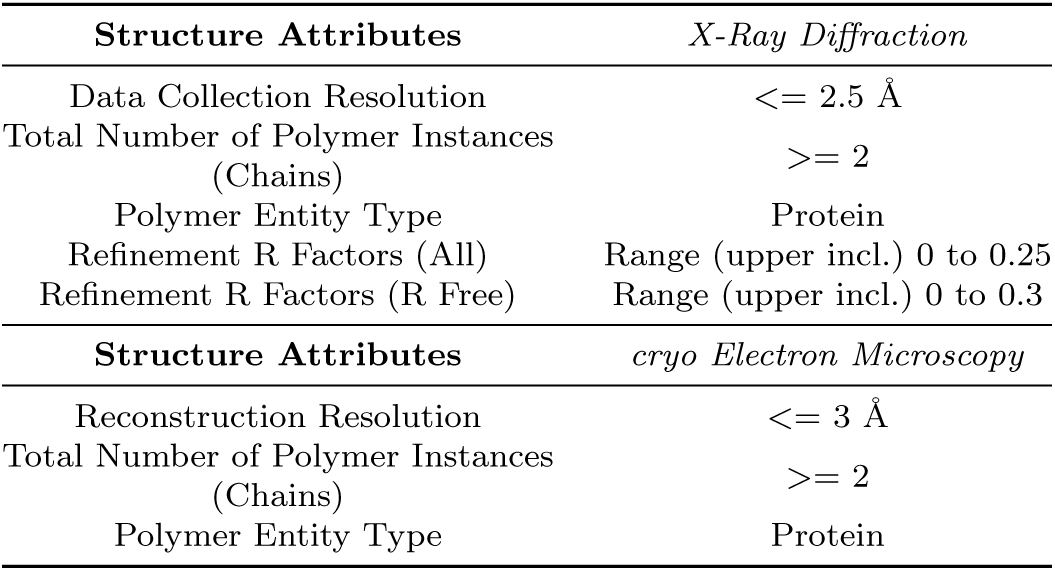
The search criteria for obtaining the positive training dataset from the Protein Data Bank.

Using these filters, 5769 X-Ray and 1916 EM resolved complexes were fetched, which were then filtered to remove all complexes with more than 30 chains, non-peptide chains and chains of under 10 residues. Upon filtering, 5005 X-Ray and 1430 EM derived complexes were retained.

Interface residues were assigned using *GEMMI* ‘s [48] distance-based neighbour search between C*α* atoms of the different chains with a cutoff of 7 Å[29]. To prevent overtraining, iAlign[49] was used to calculate the similarity among the protein-protein interfaces within a complex. The similar (symmetric) interfaces [IS-score (*>*= 0.7) and iRMSD (<= 3 Å)] within a complex were removed.

As a result, X-Ray set was reduced from 10,585 interfaces to 7092, and the EM set from 9272 interfaces to 5733. This gave a total set of 12,825 high-quality, unique, positively labeled dataset (PD1, Table 1).

#### 4.1.2 Dataset of ‘near-native’ Complexes (Positive Dataset 2 - PD2)

Protein–protein docking method *ZDOCK* [50] was used to generate ‘noise’ by creating interface poses of the interacting chains. This ‘near-native’ set comprises of the selected set of complexes with the IS-score (interface-similarity score) *>*= 0.7 and iRMSD of *<*= 3Å, which were calculated using the program iAlign[49]. This set comprises of 4006 interfaces (Table 1).

#### 4.1.3 Negative Dataset (ND)

The docked conformations with IS-score *<* 0.3 and iRMSD of *>* 4Å serve as a negative dataset, as these were structurally far from the native interfaces. This set comprises of 9090 interfaces (Table 1).

#### 4.1.4 Test Datasets

##### CASP oligomeric cryoEM targets

In order to evaluate the performance of the CNN model, oligomeric cryoEM targets from the two most recent CASP rounds, CASP15 and CASP16 were used (see Table 1 and Supplementary Table 1). For each of the targets in the CASP experiments, a pool of submitted models and an experimentally solved structure (target) are available from the CASP website.

##### PowerFit models

cryoEM structures with resolution worse than 3 Å were used. The maps were filtered by Gaussian convolution using UCSF ChimeraX[42] with a standard deviation of 2 sigma. This blurs the map by attenuating high-resolution information and gives room for rigid body fitting to generate solutions with misplacements of individual subunits. The structure is split into it’s chains and the combinations of chains with interfaces are then ran through PowerFit[35] to generate multiple fitting solution.

Interfaces from both CASP and PowerFit models with IS-score *>* 0.7 and iRMSD < 3 Å with target structure were assigned positive labels, whereas those with IS-score *<* 0.3 and iRMSD *>* 4 Å were assigned negative labels labels to the interfaces, hereafter named ‘labelled interfaces’ (Table 1). Interfaces that fall outside these thresholds are referred to as unlabelled interfaces.

### 4.2 3D CNN Network Architecture

An overview of the 3D CNN model architecture used is shown in Fig. 1a. The input is a 3D volumetric grid that represents the X, Y and Z coordinates of each atom type in the interface residues: carbon (C), nitrogen (N), oxygen (O), and sulfur (S). For computational efficiency, the input grids were down-sampled by a factor of four through a 3D pooling operation after convolutional layer 1 and 2 (Conv1 and Conv2), thus for an input grid size of 32, the total receptive field at the output features is 8 voxels. To enhance stability in feature learning, a residual skip connection is incorporated at the final convolutional layer. The extracted grid features are aggregated over the channel dimensions through global pooling and passed to the fully connected network, which produces an output of shape (N,2). This output is subsequently passed through a Soft-max layer to generate the probabilities of the interface belonging to the positive (close to target/native interface, Class 1) or negative (far from target/native interface, Class 0) interface.

### 4.3 Dynamic Grid Scaling

The ‘simple grid’ representation maps the coordinates of each atom to a voxel cell of a predefined fixed grid size *M* ^3^ using the nearest Cartesian coordinates. Although straightforward, this approach is prone to truncated or quantised representations as multiple atoms can be mapped to a single voxel, resulting in the loss of atomic information.

In ‘dynamic grid’ representation, a 3D voxel grid at 1 Å^3^ resolution covering the interface residues is created at first and the atomic coordinates to individual voxels according to nearest Euclidean distance from the voxel centre are mapped. The resulting grid is then volumetrically rescaled to the standardised dimensions *M* ^3^ as required for the downstream tasks.

The predefined fixed grid size in simple grid can be increased to reduce the atomic overlapping, particularly for large interfaces, however, this substantially increases the computational burden in the downstream tasks (see Fig. 1d). For instance, increasing the grid size to 48^3^ from 32^3^ (a 50% increase) results in approximately 3.5 fold longer epoch training times. To overcome this limitation, we resorted to a ‘dynamic’ grid representation approach [51, 52] where the grid dimensions *H* × *D* × *W* are determined adaptively based on the size of the interface.

### 4.4 Experimental Settings

All models were trained using AdamW[53] and binary cross entropy (BCE) loss using a Pytorch framework[54] with batch size of 32 and an initial learning rate of 0.0003. The learning rate is reduced by a factor of 0.95 if validation loss doesn’t improve for five epochs. The initial number of epochs for training was set to to 100 with early stopping employed if the validation loss failed to improve for five epochs.

Weight decay with cosine annealing scheduler that progressively increases the regularisation strength from 0.05 at the start of training to 0.5 by the final epoch was used. Additionally, dropout with a probability of p=0.2 is applied to the output of each convolutional layer to reduce redundant features and reliance on specific activation pathways. The purpose of these regularization techniques is to prevent the model from over-fitting.

### 4.5 K-Fold Cross Validation

The complete PDB dataset was partitioned into five non-overlapping folds using stratified sampling to preserve class distribution across fold. Five-fold cross-validation was performed, where in each fold one subset (20%) was used as the validation set and the remaining four subsets (80%) were used for training. Consequently, for each DPI model variant, five independently trained models were obtained (one per fold) and each model was evaluated on its corresponding validation fold using the performance metrics (see section 4.9).

As the training datasets are non-redundant representatives at 50% sequence identity, we didn’t explicitly enforce homolog groups using structural similarity during cross-validation. Nevertheless, we quantified cross-fold interface similarity using pair-wise iAlign comparisons, which showed that the proportion of similar interfaces between any two folds is consistently low (*<* 0.06%) indicating minimal structural overlap across folds (see Supplementary Table 5).

### 4.6 Data Augmentation

In the model with data augmentation(DPI-IV), random rotations between [−60°, 60°] and random scaling by a factor of [0.95, 1.1] were applied. These transformations help the model become invariant to minor spatial variations and orientation-dependent features, which are common in protein interfaces due to natural conformational flexibility and structural diversity[30, 55].

### 4.7 Continuous Atom Encoding

The discrete encoding of the atom types in the voxel grids (0 or 1) may limit the model’s ability to learn distinct features from the protein-protein interface. To address this, and following previous works[56, 57], atom type was represented as a density distributions around the atom centre. More specifically, at each voxel grid containing an atom, 3^3^ Gaussian kernel was applied with centre at the voxel’s origin and standard deviation set to the van der Waals radius of the corresponding atom type. The model trained with such grid representations converged more consistently and exhibited higher accuracy than the discrete representations.

### 4.8 Comparison with other scoring functions

The performance of the DPI-score was compared with the two recent state-of-the art methods for scoring protein-protein interfaces, namely PI-score[29] and DeepRank-GNN-ESM[32]. The cutoff used for PI-score for assigning true labels were: PI-score *>* 0: positive and PI-score *<*= 0: negative. The output score of DeepRank-GNN-ESM is a predicted fnat (fraction of native contacts), interfaces with predicted fnat of *>*= 0.1 were classified as positive and interfaces with *<* 0.1 were classified as negative interfaces.

### 4.9 Performance Metrics

A brief summary of fundamental metrics used in the classification tasks - Accuracy, Precision, Recall, and F1-Score - is listed below.

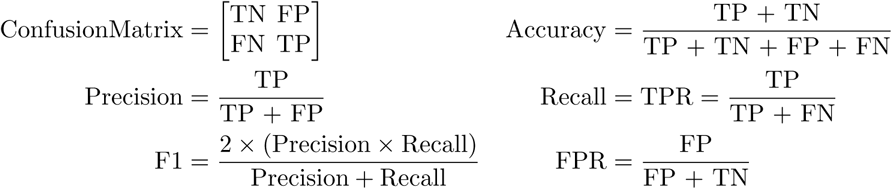

where;

y_true is the ground truth class label (1 for PD1 and PD2, 0 for ND).

y_pred is the predicted class label (1 for PD1 and PD2, 0 for ND).

True Positive (TP) is number of correctly classified positive instances (y_true=1, y_pred=1).

True Negative (TN) is number of correctly classified negative instances (y_true=0, y pred=0).

False Positive (FP) is number of negative instances incorrectly classified as positive (y_true=0, y_pred=1).

False Negative (FN): Positive instances incorrectly classified as negative (y_true=1, y_pred=0).

Beside these metrics we also report single threshold independent metrics like ROC curves (TPR vs FPR), PR curves (precision vs recall) and a single ROC-AUC metric were reported to help estimate the ranking capabilities of DPI-score.

### 4.10 Density-based scores

MolProbity[22] evaluates different stereochemical aspects of the model like the quality of bonds, angles and dihedrals and serious atomic clashes (clashscore). The MolProbity score is a global estimate of model quality derived using a weighted combination of selected individual geometry scores. Lower values of the MolProbity score reflect better geometry. Clashscore reflects the overall severity of atomic clashes in the model and lower values indicate fewer severe clashes.

Average model-map FSC 0.5 score is derived from the model–map FSC curve by calculating an average of the FSC weighted by the number of structure factors in each shell[58] up to the resolution corresponding to FSC=0.5. A cross correlation coefficient (CCC) can also be calculated in real-space between the model and the map to estimate the quality of model fit. To mimise the effect of background and unmodelled regions, this can be calculated locally within the region of overlap between the model and map (CCC mask).

The segment-based Manders’ overlap coefficient (SMOC) score quantifies the per-residue agreement between a theoretical map derived from the atomic model and the experimental map[20]. To identify outliers or potential misfits, we compute a Z-score for each residue relative to the local neighborhood (residues within 12 Å). SMOC Z-score *<* −2 gives the fraction of residues with Z-scores *<* −2, suggesting that the fit could potentially be improved. Lower SMOC Z-score *<* −2 values indicate fewer such residues.

### 4.11 Dataset Biases

The model was trained on experimentally determined structures from the Protein Data Bank (PDB) and evaluated on PowerFit Models, CASP targets, and EMDB benchmarks. The training datasets are non-redundant representatives at 50% sequence identity. The pairwise interface similarity between any folds are less then 0.06% as shown in Supplementary Table 5.

## 5 Acknowledgements

We thank the CCP-EM team for useful discussions and Dr. Andriy Kryshtafovych (UC Davis) for the help with CASP targets. The authors (NB, APJ, TS and SM) would like to acknowledge support from the Ada Lovelace Centre, STFC, UKRI. SM would also like to acknowledge Artificial Intelligence in the Biosciences - AIBIO-UK (22-AIBN) [BB/Y006933/1]. APJ would also like to thank the MRC for Partnership Grant MR/V000403/1. The authors NW and Jyothish Thomas were partially supported by the BASE-II ExCALIBUR project (EPSRC grant number EP/X019918/1), which focused on identifying and developing AI for Science methodologies with the potential to scale to large-scale and exascale scientific applications.

## 6 Author Contributions

N.B., A.P.J., and S.M. designed the research. N.B., N.W., Jyothish Thomas, A.P.J. and S.M. wrote the software. T.S. performed software engineering and testing of the software. N.B, N.W., L.E. and S.M. performed the experiments, data analysis. N.B, N.W., A.P.J and S.M. wrote the first draft of the paper. N.B, N.W., A.P.J., Jeyan Thiyagalingam and S.M contributed to paper revision.

## 7 Data availability

All the structures and assessed models are deposited in the freely accessible public databases-PDB and EMDB. All the PDB IDs and EMDB IDs are appropriately cited throughout the manuscript. All the results generated during this study are within the article and its Supplementary files.

## 8 Code availability

The software to calculate the DPI-score is freely available for academic use through: GitLab at https://gitlab.com/ccpem/dpi or via the CCP-EM software suite, Doppio, at https://www.ccpem.ac.uk/software

## Supplementary Information

**Supplementary Figure 1:**
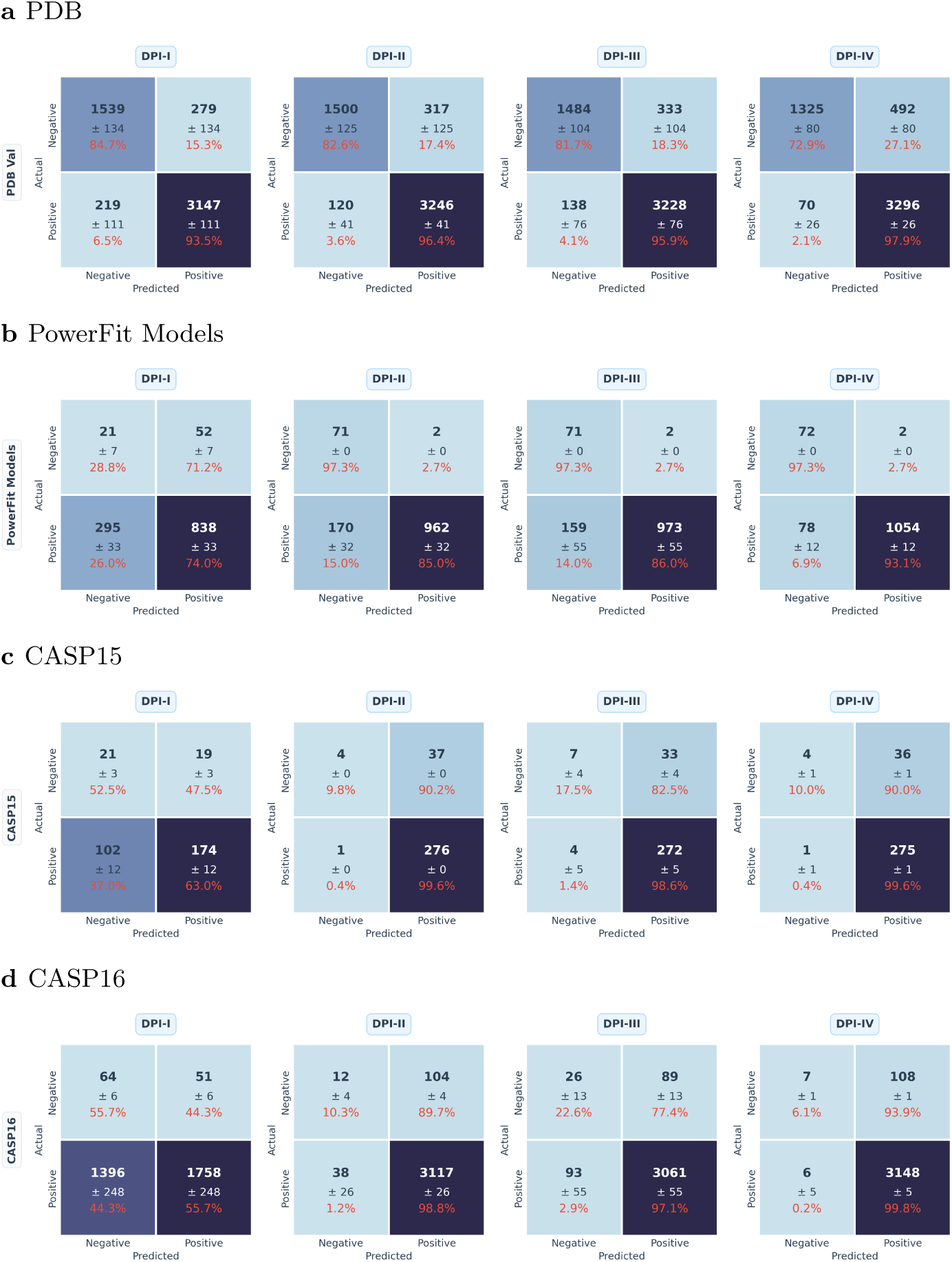
Confusion matrices aggregated over five folds of the four DPI variants across test datasets.

**Supplementary Figure 2:**
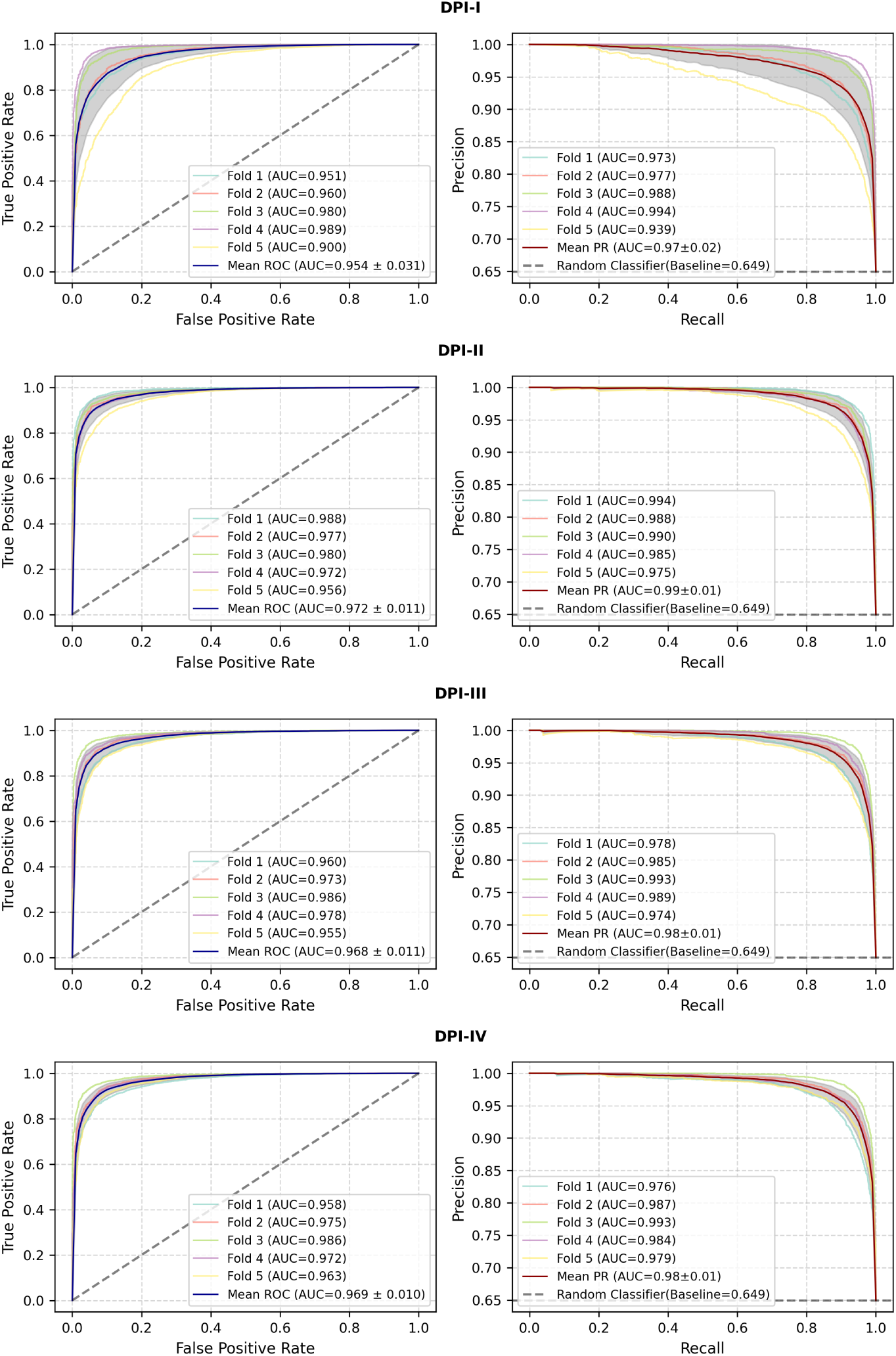
Receiver Operating Characteristic (ROC) curve (left) and Precision-Recall curve (right) of all DPI models on 5-fold PDB validation set.

**Supplementary Figure 3:**
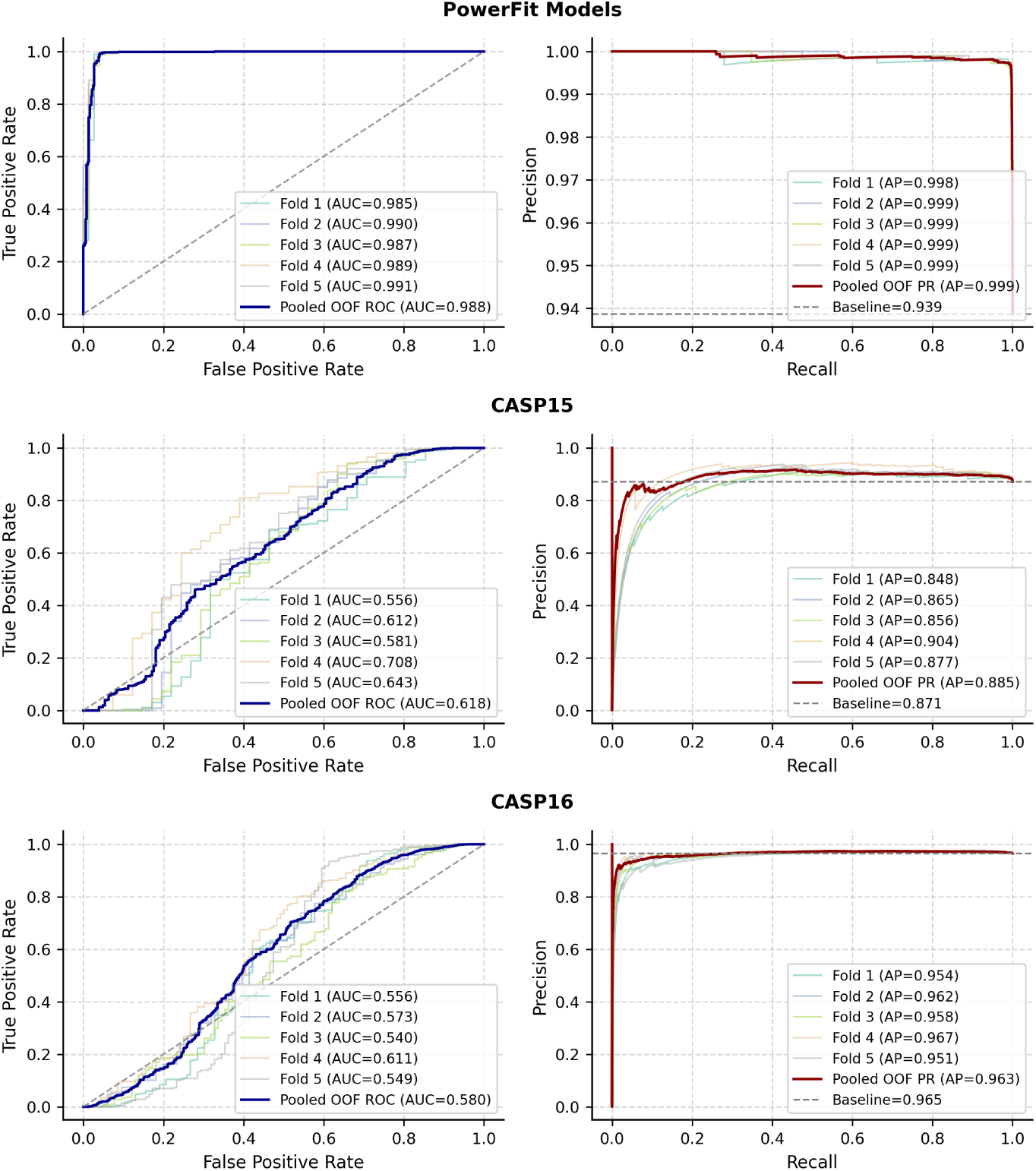
ROC curves (left) and Precision-Recall curve (right) of DPI-IV on test datasets.

**Supplementary Figure 4:**
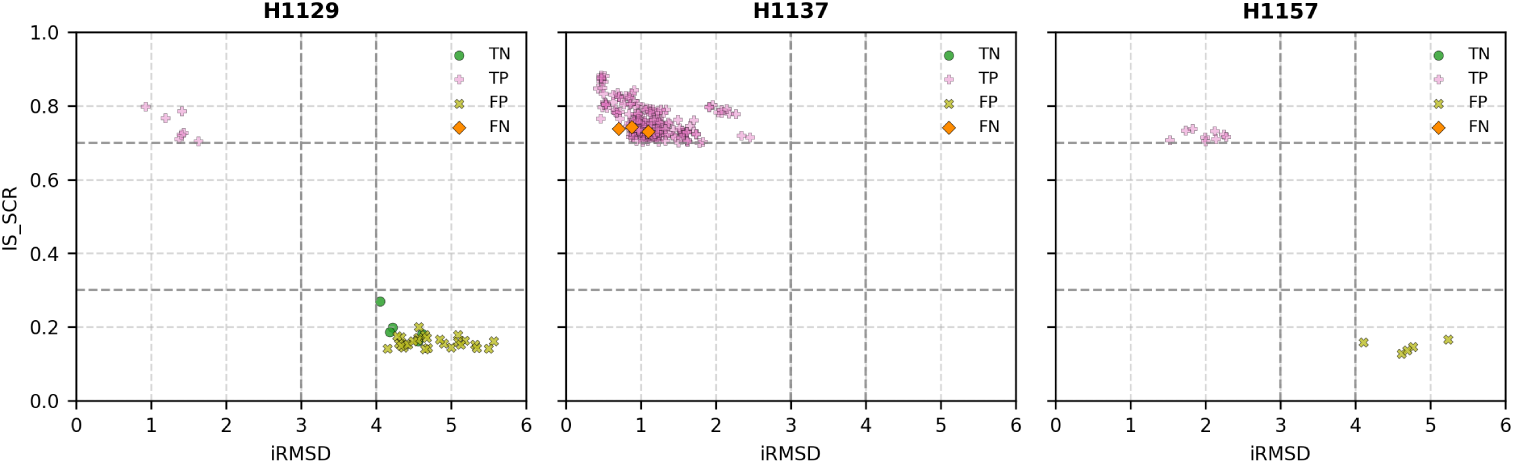
IS-Score vs iRMSD plot of labelled interfaces in CASP15 targets and corresponding predictions (TP, TN, FP and FN) from DPI-IV. Dashed lines indicates the threshold on IS-score and iRMSD for interface labelling.

**Supplementary Figure 5:**
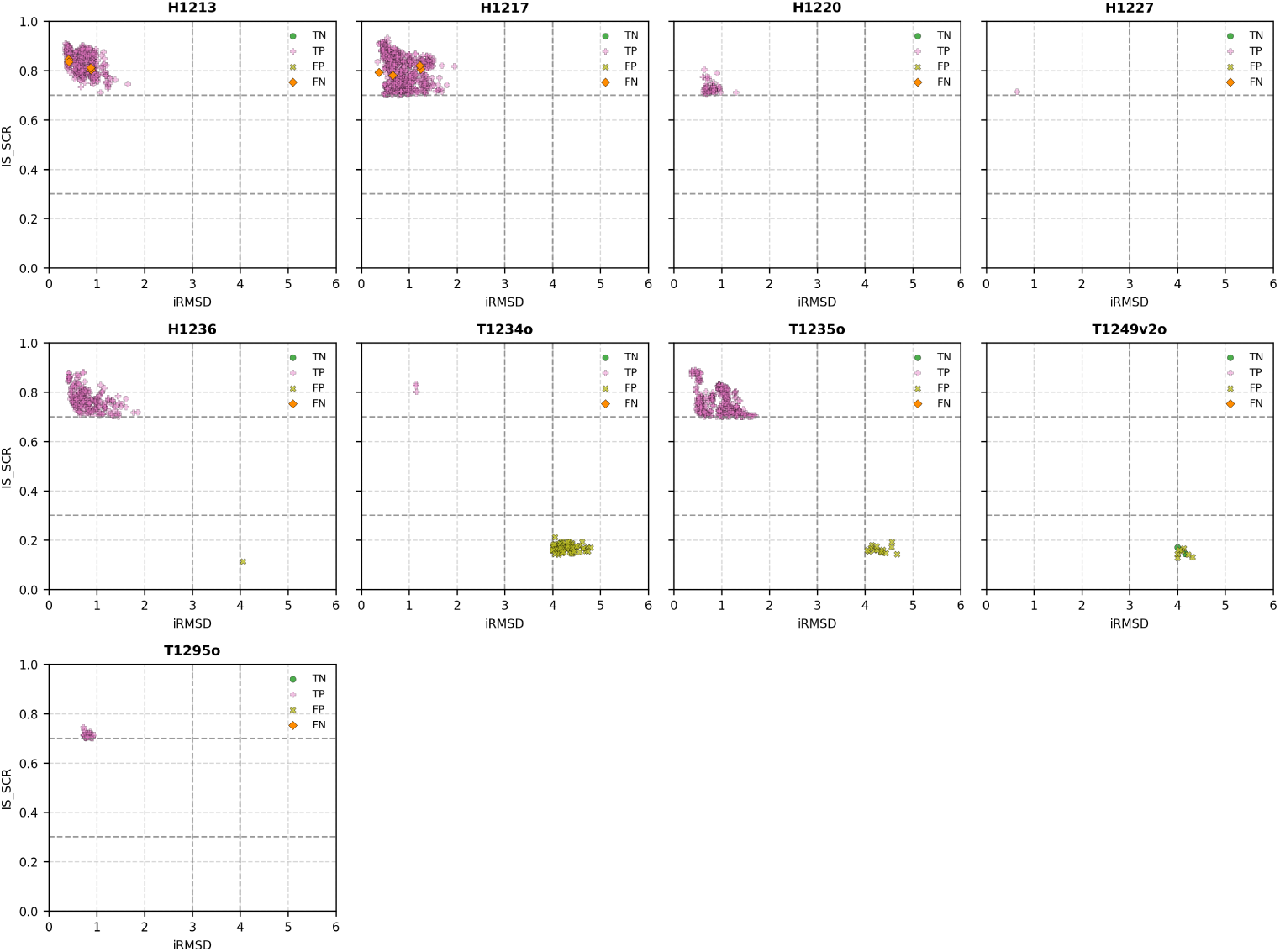
IS-Score vs iRMSD plot of labelled interfaces in CASP16 targets and corresponding predictions (TP, TN, FP and FN) from DPI-IV. Dashed lines indicates the threshold on IS-score and iRMSD for interface labelling.

**Supplementary Figure 6:**
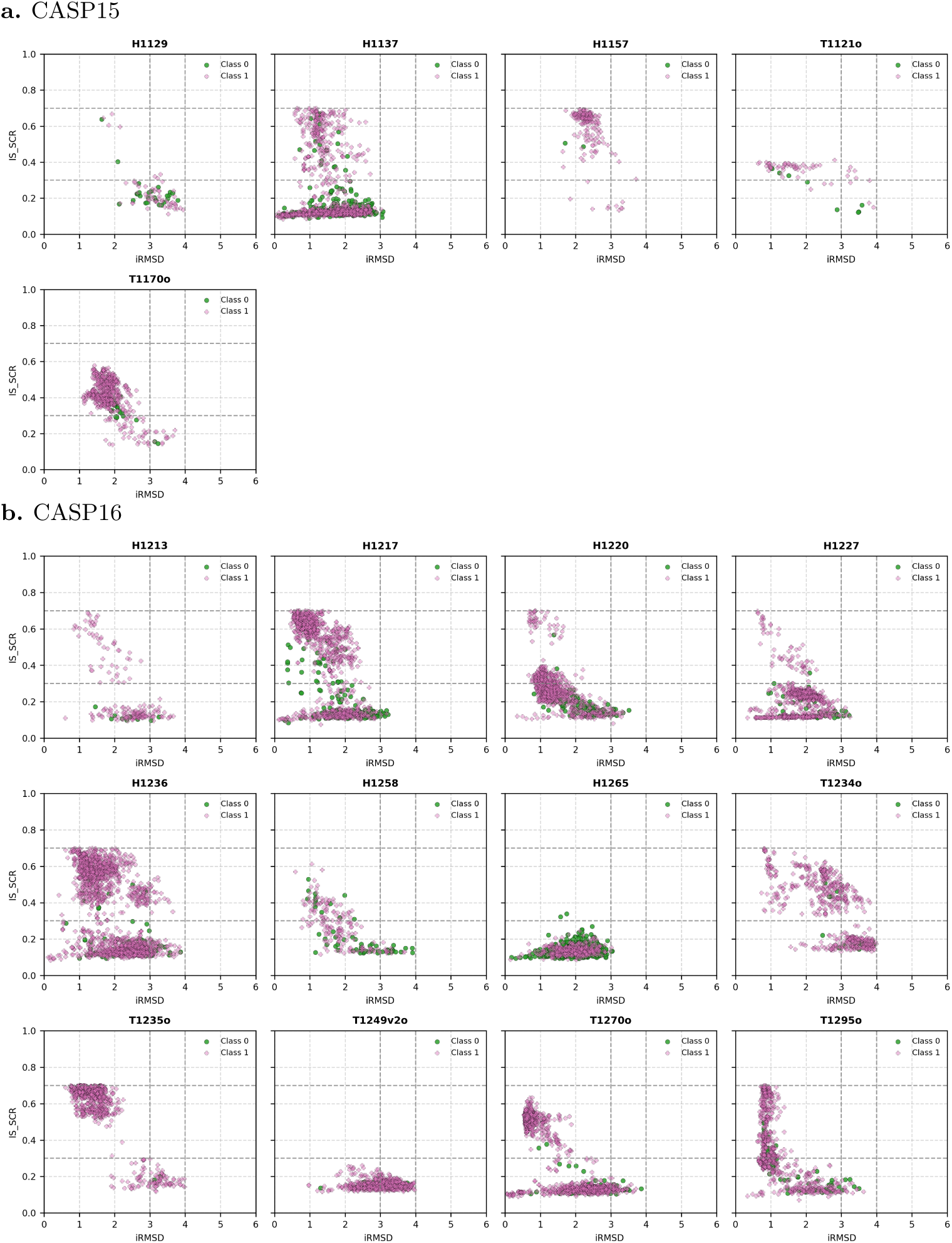
IS-Score vs iRMSD plot for unlabelled interfaces **a**. CASP15 targets, **b**. CASP16 targets. Class 0 represents interfaces predicted to have lower DPI-scores (*<* 0.5) and Class 1 represents interfaces with higher DPI-scores (*>*= 0.5). Dashed horizontal and vertical lines shows the threshold values used for IS-score and iRMSD respectively for interface assignment.

**Supplementary Figure 7:**
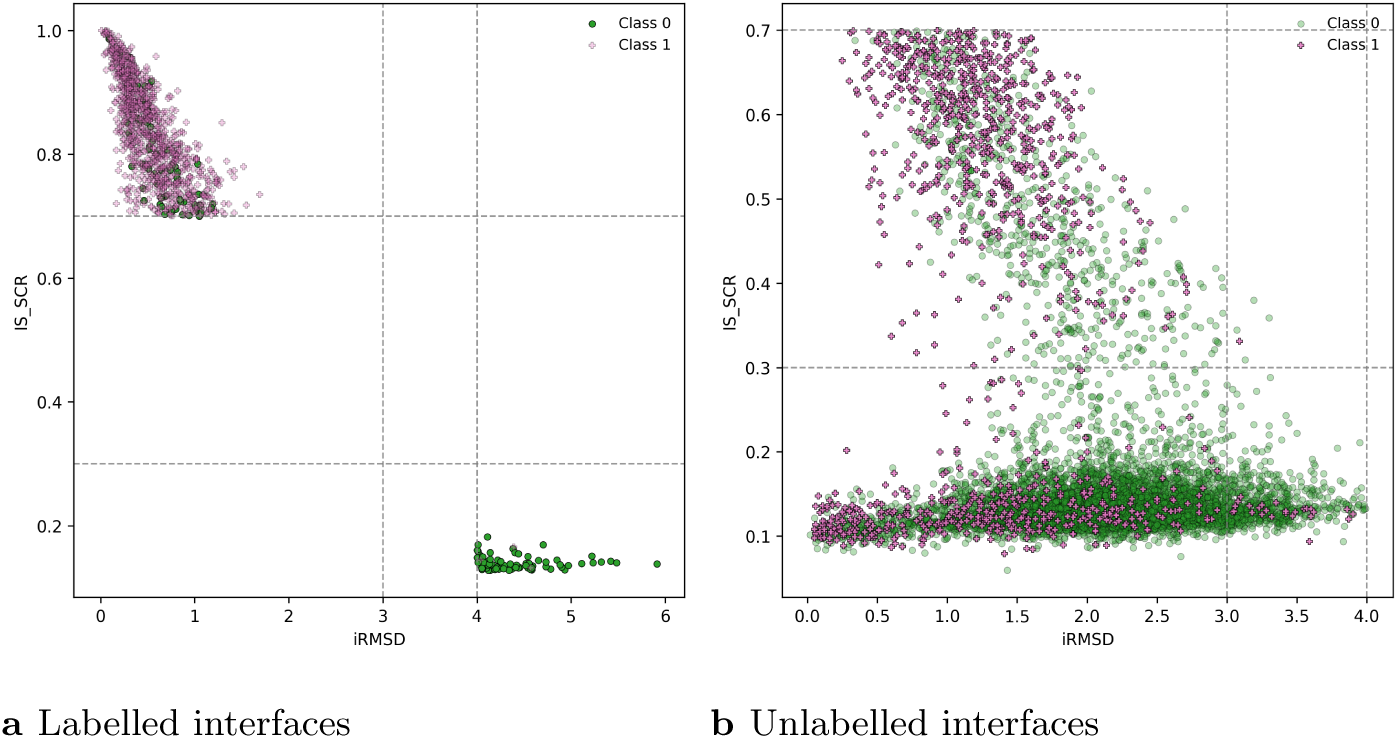
IS-Score vs iRMSD plot of interfaces from PowerFitted models. Class 0 represents interfaces predicted to have lower DPI-scores (*<* 0.5) and Class 1 represents interfaces with higher DPI-scores (*>*= 0.5). Dashed lines indicates IS-score and iRMSD thresholds used for interface labelling.

**Supplementary Figure 8:**
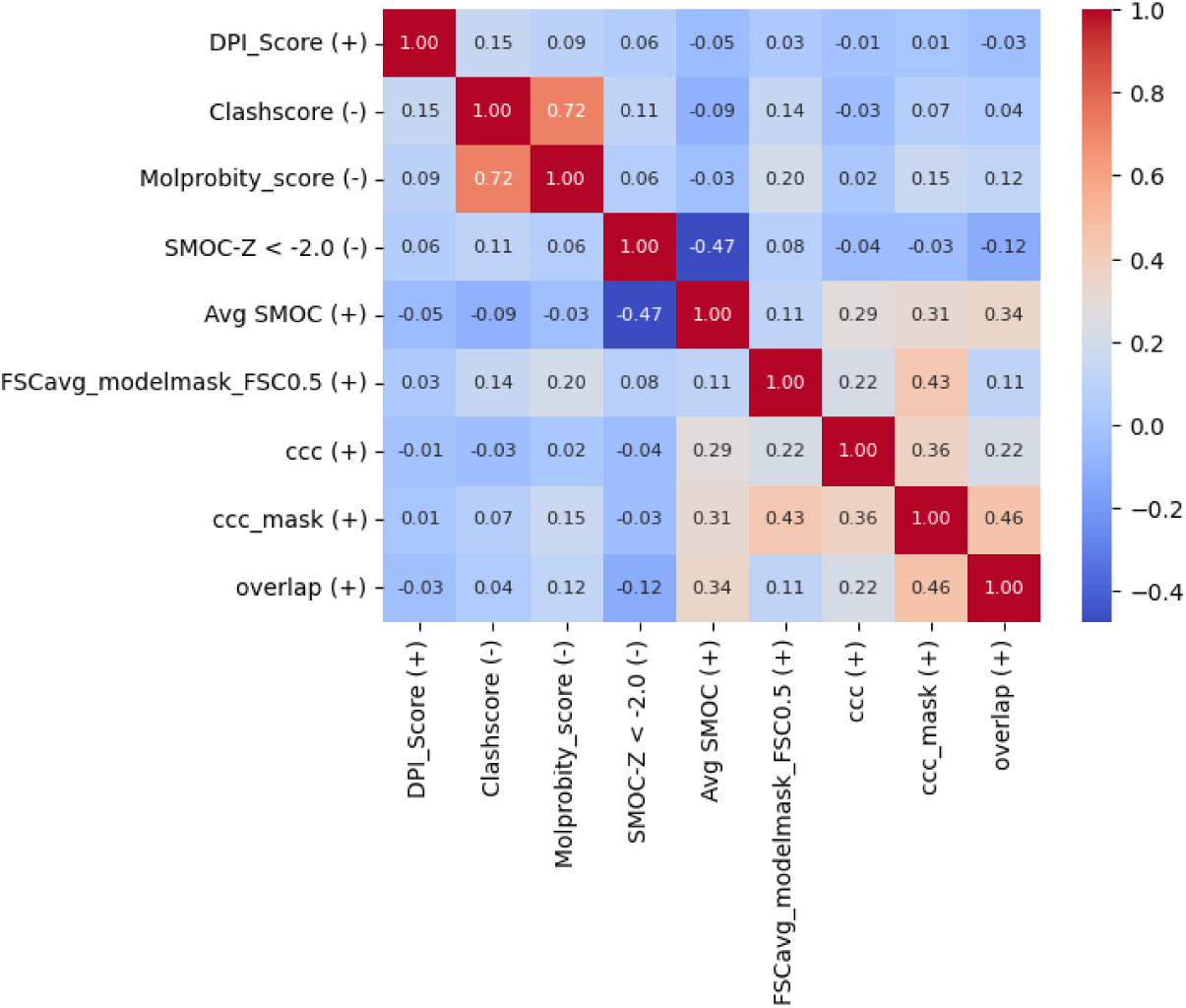
Correlation of DPI-score with density-based scores. (+) indicates higher is better, (–) indicates lower is better.

**Supplementary Figure 9:**
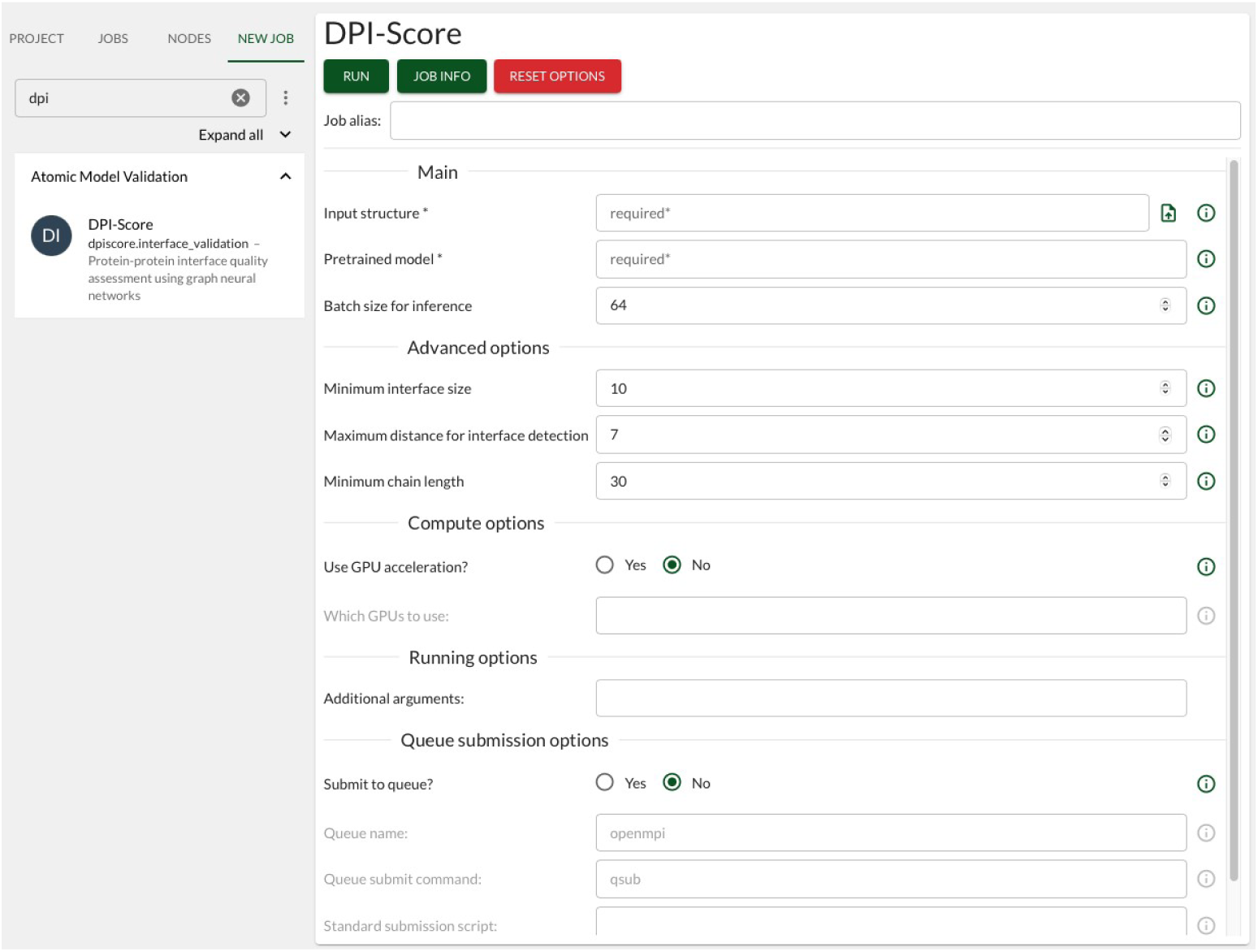
A screenshot of Doppio interface to run DPI-Score. Doppio is a free (for non-profit use) web-based user interface that pipeline multiple validation tasks in a single interface. Please see https://www.ccpem.ac.uk/docs/doppio for installation instructions.

**Supplementary Table 1:**
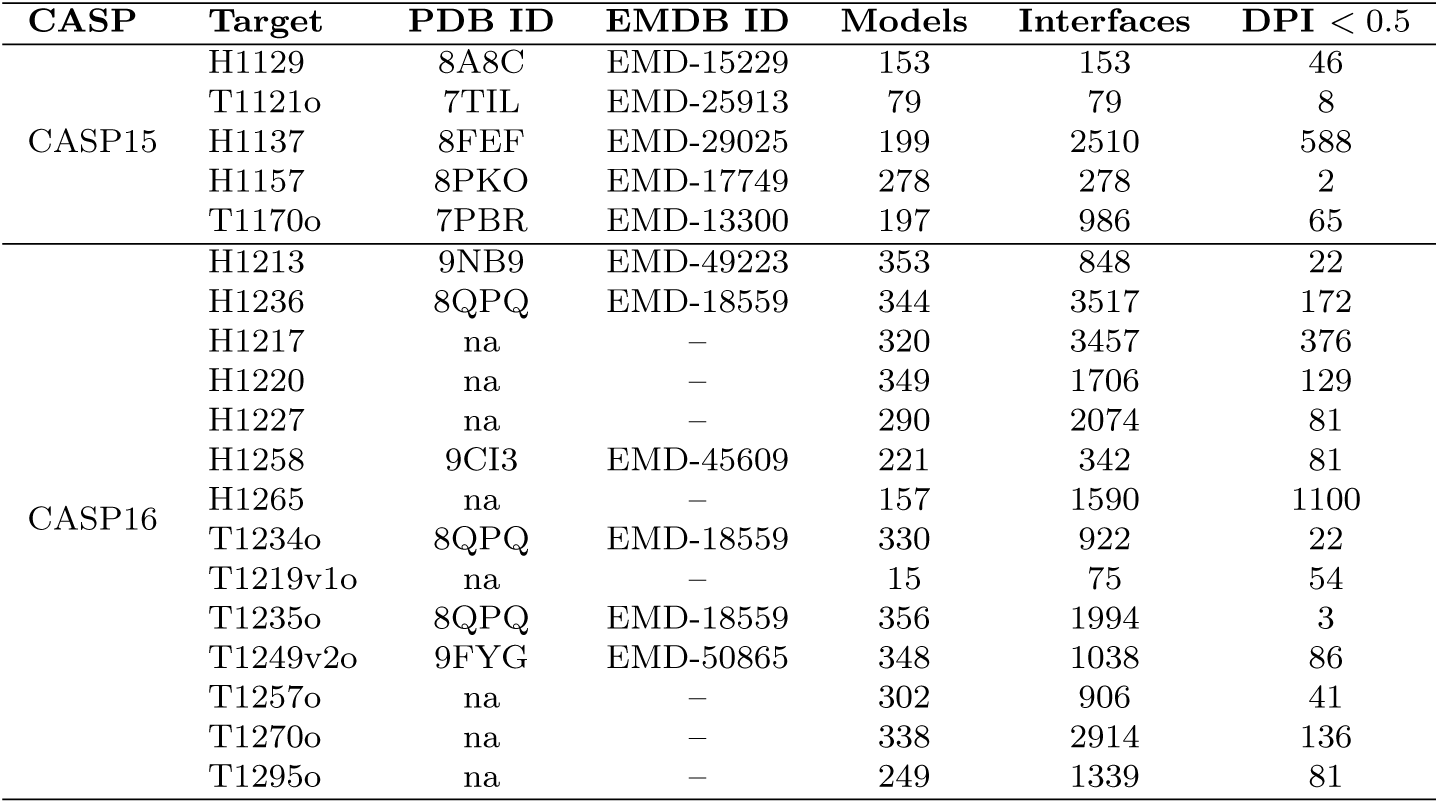
Oligomeric cryoEM targets from CASP15 and CASP16 used for testing. PDB ID and EMDB ID are listed where available. na indicates that the structure was not released at the time of analysis. The total number of deposited models, assessed interfaces, and interfaces with DPI-score *<* 0.5 are shown.

| CASP | Target | PDB ID | EMDB ID | Models | Interfaces | DPI $< 0.5$ |
| --- | --- | --- | --- | --- | --- | --- |
| CASP15 | H1129 | 8A8C | EMD-15229 | 153 | 153 | 46 |
|  | T1121o | 7TIL | EMD-25913 | 79 | 79 | 8 |
|  | H1137 | 8FEF | EMD-29025 | 199 | 2510 | 588 |
|  | H1157 | 8PKO | EMD-17749 | 278 | 278 | 2 |
|  | T1170o | 7PBR | EMD-13300 | 197 | 986 | 65 |
| CASP16 | H1213 | 9NB9 | EMD-49223 | 353 | 848 | 22 |
|  | H1236 | 8QPQ | EMD-18559 | 344 | 3517 | 172 |
|  | H1217 | na | — | 320 | 3457 | 376 |
|  | H1220 | na | — | 349 | 1706 | 129 |
|  | H1227 | na | — | 290 | 2074 | 81 |
|  | H1258 | 9CI3 | EMD-45609 | 221 | 342 | 81 |
|  | H1265 | na | — | 157 | 1590 | 1100 |
|  | T1234o | 8QPQ | EMD-18559 | 330 | 922 | 22 |
|  | T1219v1o | na | — | 15 | 75 | 54 |
|  | T1235o | 8QPQ | EMD-18559 | 356 | 1994 | 3 |
|  | T1249v2o | 9FYG | EMD-50865 | 348 | 1038 | 86 |
|  | T1257o | na | — | 302 | 906 | 41 |
|  | T1270o | na | — | 338 | 2914 | 136 |
|  | T1295o | na | — | 249 | 1339 | 81 |

**Supplementary Table 3:** Cross-correlation coefficient (CCC) values for T1121o models at lower resolution values.

| Model | DPI | CCC(3.7 Å) | CCC(5 Å) | CCC(8 Å) | CCC(10 Å) | CCC(12 Å) |
| --- | --- | --- | --- | --- | --- | --- |
| TS098.5o_A_B | 0.03 | 0.62 | 0.65 | 0.66 | 0.77 | 0.82 |
| TS165.2o_A_B | 0.20 | 0.55 | 0.57 | 0.62 | 0.64 | 0.66 |
| TS098.4o_A_B | 0.32 | 0.59 | 0.61 | 0.66 | 0.67 | 0.69 |
| TS234.1o_A_B | 0.36 | 0.47 | 0.50 | 0.56 | 0.58 | 0.61 |
| 7TIL (target) | 0.99 | 0.84 | 0.89 | 0.94 | 0.95 | 0.96 |

**Supplementary Table 5:** Percentage of similar pairs between folds. The percentage are calculated as ratio of number of pairwise interfaces with iAlign score *>*= 0.7 to the total number of pairwise interfaces in each fold pair.

| **Fold** | **2** | **3** | **4** | **5** |
| --- | --- | --- | --- | --- |
| **1** | 0.052% | 0.054% | 0.059% | 0.057% |
| **2** |  | 0.050% | 0.057% | 0.046% |
| **3** |  |  | 0.060% | 0.048% |
| **4** |  |  |  | 0.054% |

## Notes

https://gitlab.com/ccpem/dpi

### Competing Interest Statement

The authors have declared no competing interest.

### Summary of Updates

Some sections from Results are moved to Methodology Include comparisons results and benchmarking with existing methods

